# DMT1 bridges endosomes and mitochondria to modulate mitochondrial iron translocation

**DOI:** 10.1101/2022.04.14.488402

**Authors:** Jonathan Barra, Isaiah Crosbourne, Ling Wang, Ramon Bossardi-Ramos, Frances Jourd’heuil, Iram Nelson, Alejandro P. Adam, David T. Corr, David Jourd’heuil, Margarida Barroso

## Abstract

Transient “kiss-and-run” endosome-mitochondria interactions can mediate mitochondrial iron translocation (MIT) but the associated mechanisms are still elusive. We show that Divalent Metal Transporter 1 (DMT1) modulates MIT via endosome-mitochondria interactions in invasive MDA-MB-231, but not in non-invasive T47D breast cancer cells. CRISPR/Cas9-based DMT1 knockout (KO) stable cells were used to demonstrate that DMT1 regulates MIT, endosomal speed, and labile iron pool (LIP) levels only in MDA-MB-231. DMT1 silencing increases PINK1/Parkin mitophagy markers, the autophagy marker LC3B, as well as mitochondrial ferritin in MDA-MB-231, but not in T47D. Strikingly, re-expression of DMT1 in MDA-MB-231 DMT KO cells rescues all protein levels evaluated. DMT1 silencing decreases Tom20 colocalization with PMPCB, a DMT1 interactor that regulates mitophagy hyperactivation. In MDA-MB-231 both mitochondrial metabolism and invasion were impaired by DMT1 silencing and rescued by DMT1 re-expression. DMT1 acts as a bridge between endosomes and mitochondria to support higher MIT/lower LIP levels, which are necessary for sustaining mitochondrial bioenergetics and invasive cancer cell migration.

**Summary:** Cellular iron metabolism is tightly regulated, and cancer cells rely on mitochondrial iron for malignancy. Here, we report that the divalent metal transporter DMT1 serves as a bridge between endosomes and mitochondria regulating mitochondrial iron translocation in breast cancer cells.

## Introduction

Iron is an essential component for several key proteins and enzymes that are required for proper cellular functioning (Andrews and Schmidt, 2007). Evolutionarily, iron requirements constitute an important mechanism for genetic selection (Wade et al., 2021). Moreover, iron homeostasis is tightly regulated to limit intracellular labile iron pool (LIP), which is associated with the generation of harmful reactive oxygen species (ROS) (Dixon and Stockwell, 2014).

Common or classical hallmarks of cancer have been extensively described (Hanahan, 2022). However, given cancer heterogeneity and complexity, other non-classical cancer hallmarks have been proposed, e.g., the iron-seeking phenotype of cancer cells (Torti et al., 2018). Epidemiologically, iron is a risk factor for different types of cancer (Fonseca-Nunes et al., 2014). In breast cancer, recent studies have suggested an association between reduced risk of breast cancer and lower levels of iron, particularly in obese women, although further studies are necessary to corroborate or discard this association (Chang et al., 2019; Von Holle et al., 2021). Furthermore, mitochondrial iron accumulation seems to rewire iron metabolism homeostasis in cancer cells towards adaptation to malignant features, including enhanced invasion and metastatic capacity (Torti and Torti, 2020).

Uptake of dietary iron is performed by enterocytes mainly via divalent metal transporter 1 (DMT1)-isoform I, which is located at the plasma membrane (Garrick et al., 2003; Gunshin et al., 2005; Hubert and Hentze, 2002). Ferric iron is then reduced to its ferrous form by duodenal cytochrome B (McKie et al., 2001), and then transported to the basolateral membrane of the enterocyte where it is exported to the circulation via ferroportin (FPN), the only known mammalian iron exporter (Drakesmith et al., 2015). Once in the circulation, ferric iron is mainly transported associated to transferrin (Tf). On the other hand, non-transferrin-bound ferric iron can be rapidly metabolized from the plasma by the liver (Craven et al., 1987) or taken up by specific cell types via Tf-independent mechanisms, such as the Zip14 transporter (Liuzzi et al., 2006). Iron-bound Tf can be internalized by cells via transferrin receptor (TfR) mediated endocytosis (Harding et al., 1983; Mayle et al., 2012). Once internalized by peripheral cells, ferric iron is readily sorted to endosomes, where it is released from Tf upon exposure to the lower pH of the endosomal lumen. Then, ferric iron is reduced by the six-transmembrane epithelial antigen of the prostate 3 (Steap3) ferrireductase, and exported from endosomes to the cytosol by DMT1 isoform II (Fleming et al., 1998; Ohgami et al., 2005; Tabuchi et al., 2010). In the cytoplasm, iron can be either bound to specific chaperones for transport to distinct cellular compartments, stored in cytosolic ferritin, or exported via FPN (Yanatori and Kishi, 2019). Alternatively, transient interactions (“kiss-and-run”) between early endosomes (EE) and mitochondria have been shown to facilitate the direct translocation of iron into the mitochondria, bypassing the cytosol (Das et al., 2016; Hamdi et al., 2016; Sheftel et al., 2007). As mentioned, free ferrous iron in the cytoplasm is cytotoxic since it induces ROS production (Jomova and Valko, 2011). Mitochondria, as the main cellular iron sink, can also store iron in mitochondrial ferritin (FTMT) for its use in heme and iron sulfur clusters biogenesis, which are components of molecules such as hemoglobin and respiratory complexes I-III, respectively (Dev and Babitt, 2017).

DMT1 (also known as SLC11A2 or NRAMP2/DCT1) is comprised of 12 transmembrane domains with *N*-glycosylation sites at C-and N-terminal tails, both facing the cytosol (Gunshin, 1997). Besides iron, DMT1 can also transport other divalent metals including Mn, Co, and Cu (Gunshin, 1997), although with less molecular affinity. Alternative splicing generates two isoforms of DMT1 mRNA, DMT1-I and DMT1-II. After proper folding in the ER and Golgi apparatus, where they are N-linked glycosylated, each DMT1 isoform follows different sorting pathways. DMT1-isoform I is located mainly at the plasma membrane in duodenal enterocytes (Garrick et al., 2003; Gunshin et al., 2005; Hubert and Hentze, 2002), whereas, DMT1-isoform II can be found at early/recycling endosomal compartment in non-epithelial cells (Fleming et al., 1998; Ohgami et al., 2005; Tabuchi et al., 2010). Structurally, the C-terminal domain of DMT1-II (henceforth referred to as DMT1 herein) contains a retromer complex binding site, allowing it to localize at the early/recycling endosomal compartment (Lucas et al., 2016; Tabuchi et al., 2010).

Interestingly, in addition to endosomes, DMT1 has been found at the outer mitochondrial membrane (OMM) (Wolff et al., 2014), suggesting its involvement in mitochondrial iron translocation (MIT). Moreover, DMT1 overexpression has been shown to increase mitochondrial iron uptake (Wolff et al., 2018). However, the exact role of DMT1 in the mechanisms regulating endosome-mitochondria interactions and MIT are still not elucidated.

In the present work, we first evaluated the role of endosomal DMT1 in transient “kiss-and-run” endosome-mitochondria interactions, mitochondrial iron translocation (MIT), and cytoplasmic iron regulation in two different breast cancer cell lines. We showed that DMT1 can be present in endosomes as well as in association with mitochondria outer membrane (OMM), acting as a bridge between both organelles. Interestingly, these DMT1-mediated bridging events are more frequent in invasive MDA-MB-231 than in non-invasive T47D breast cancer cells. In contrast to non-invasive breast cancer cells, mitophagy can be regulated by DMT1 in invasive breast cancer cells. DMT1 also regulates cytoplasmic iron levels, as well as mitochondrial ROS generation, in invasive cancer cells. We provide evidence indicating that, compared to T47D cells, MDA-MB-321 cells are highly sensitive to DMT1 silencing, which disrupts endosome-mitochondria interactions and MIT with functional consequences in invasive migration and mitochondrial bioenergetics. These findings reveal a DMT1-dependent pathway connecting endosome-mitochondria transient interactions to MIT, mitochondrial metabolism, and invasive capacity in breast cancer cells.

## Results

### DMT1 acts as a bridge between endosomes and mitochondria

Here, we studied the role of DMT1 in the endosome-mitochondria interactions in breast cancer cells. First, cells were subjected to a 2-min pulse with fluorescently labeled Tf (EE marker; green) and chased for 2-5 min prior to fixation and immunofluorescence using anti-Tom20 (magenta), a marker of the OMM, and anti-DMT1 (red). We evaluated the co-distribution of Tf, Tom20, and DMT1 in both invasive MDA-MB-231 and non-invasive T47D breast cancer cells (**Figure 1A and 1B**). After 3D rendering of each of the organelle markers, we measured the Surface Contact Area (SCA) of both Tf with DMT1 (yellow) and DMT1 with Tom20 (cyan) (**Figure 1A(g-i) and 1B(g-i)**). In MDA-MB-231 as well as in T47D, we observed that DMT1 co-distributed with Tf as well as Tom20, which is in accordance with previous reports (Lucas et al., 2016; Tabuchi et al., 2010; Wolff et al., 2018). Strikingly, in MDA-MB-231 cells, almost all of the Tf-DMT1 SCA events were observed in close proximity to DMT1-Tom20 SCA events. In contrast, the SCAs of Tf-DMT1 and DMT1-Tom20 were found mainly separated in T47D cells (**Figure 1A and 1B, see zoom-ins (j-m)**). Taken together, these data indicate that DMT1 is associated with EE and mitochondria in breast cancer cells. Moreover, the presence of DMT1 acting as a bridge between EE and mitochondria is frequently observed in invasive MDA-MB-231 cells, but not in non-invasive T47D cells.

**Figure 1.**
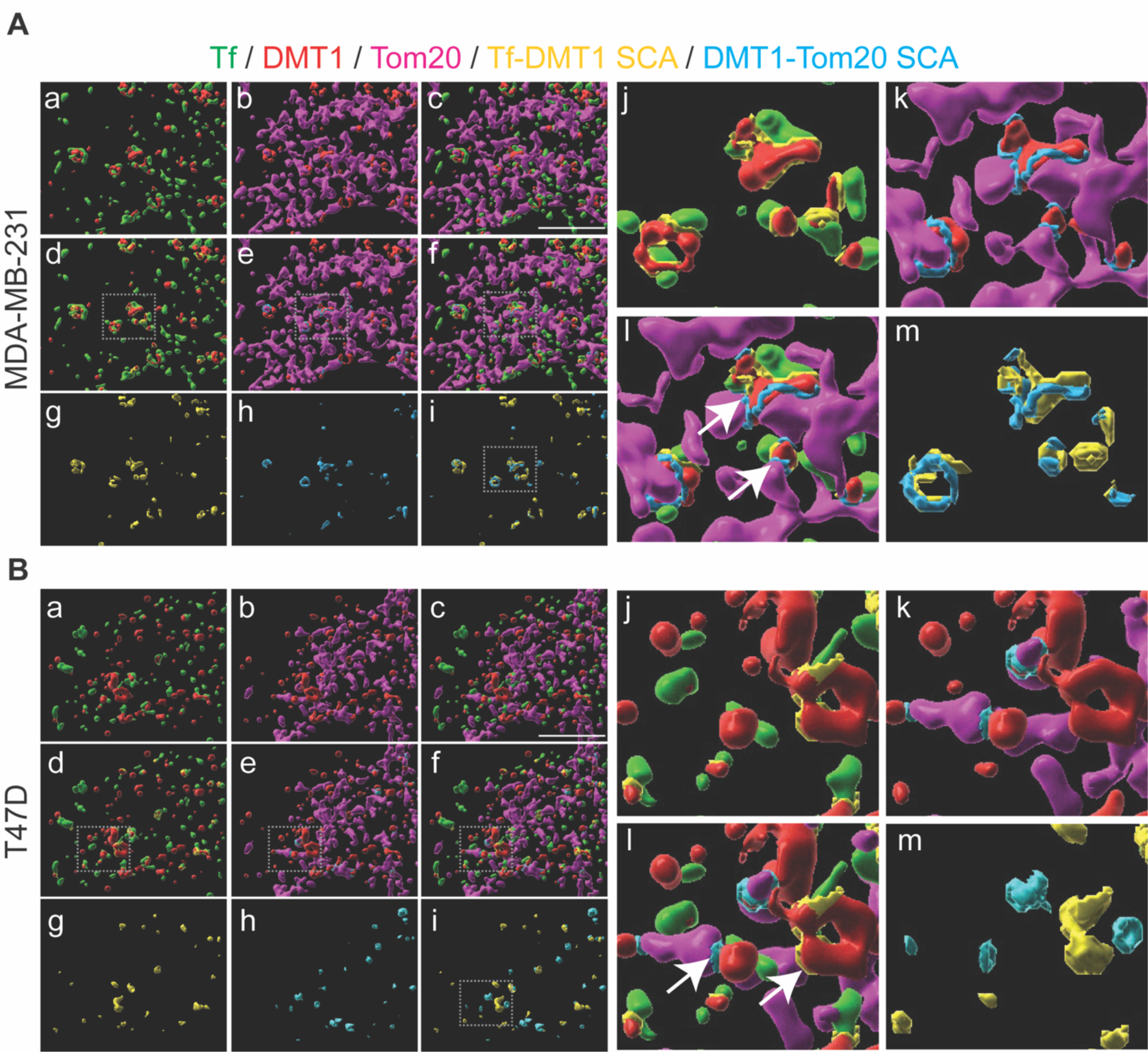
DMT1 acts as a bridge between endosomes and mitochondria in MDA-MB-231 but not in T47D breast cancer cells. **(A)** MDA-MB-231 and **(B)** T47D cells were subjected to a 2 min pulse with fluorescently labeled Tf (green) and chased for 2-5 min prior to fixation and immunofluorescence using anti-Tom20 (magenta), a marker of the OMM, and anti-DMT1 (red). Images were acquired using the Leica Thunder microscope (Objective HCX PLS-APO 63x/1.30 oil) and subjected to deconvolution using integrated SVCC algorithm. Z-stacks were 3D rendered using IMARIS software 9.6 (Bitplane) and Surface Contact Area (SCA) between Tf-DMT1 (yellow) and DMT1-Tom20 (cyan) was calculated using the “Surface Surface Contact Area” IMARIS XTension integrated plugin. **(a)** Tf and DMT1; **(b)** DMT1 and Tom20; **(c)** Tf, DMT1 and Tom20; **(d)** Tf, DMT1 and Tf-DMT1 SCA; **(e)** DMT1, Tom20 and DMT1-Tom20 SCA; **(f)** Tf, DMT1, Tom20, Tf-DMT1 SCA and DMT1-Tom20 SCA; **(g)** Tf-DMT1 SCA; **(h)** DMT1-Tom20 SCA; **(i)** Tf-DMT1 SCA and DMT1-Tom20 SCA; **(j)** magnified region from region-of-interest shown in **(d)**; **(k)** magnified region from region-of-interest shown in **(e)**; **(l)** magnified region from region-of-interest shown in **(f)**; **(m)** magnified region from region-of-interest shown in **(i)**. Dotted squares indicate the region-of-interest highlighted. Lower association between Tf-DMT1 and Tom20-DMT1 SCAs in T47D compared with MDA-MB-231 is observed. Scale bar = 10 μm.

### DMT1 regulates association between endosomes and mitochondria

In epithelial cells, we previously reported that the kinetics of “kiss-and-run” endosome-mitochondria interactions are modulated by the availability of intra-endosomal iron (Das et al., 2016). However, the mechanisms and molecules involved in this process, and how they can be regulated in cancer cells, are mainly unexplored. CRISPR/Cas9 technology was used to silence DMT1 in both MDA-MB-231 and T47D breast cancer cells (**Figure 2A**). Live-cell time-lapse imaging of cells co-labeled with Tf and Mitotracker (Mito) was used to evaluate Tf-Mito “kiss-and-run” transient interactions (**Figure 2B**) in wild-type (WT) and DMT1 knock-out (KO) cells. After 3D rendering of Tf and Mito organelle objects in live-cell imaging experiments (interval 2.5 seconds, total time 45 seconds), we measured the sum of the SCA between Tf- and Mito-labeled objects as an indicator of the total EE-mitochondria interactions per cell. We found that the sum of Tf-Mito SCA per cell decreased significantly upon DMT1 silencing in MDA-MB-231 but not in T47D cells (**Figure 2C**). Although the average of Tf-Mito SCA per cell was similar between WT and DMT1 KO in both cell lines (**Figure 2D**), a significant decrease was observed in T47D WT compared to MDA-MB-231 WT (**Figure 2D**).

**Figure 2.**
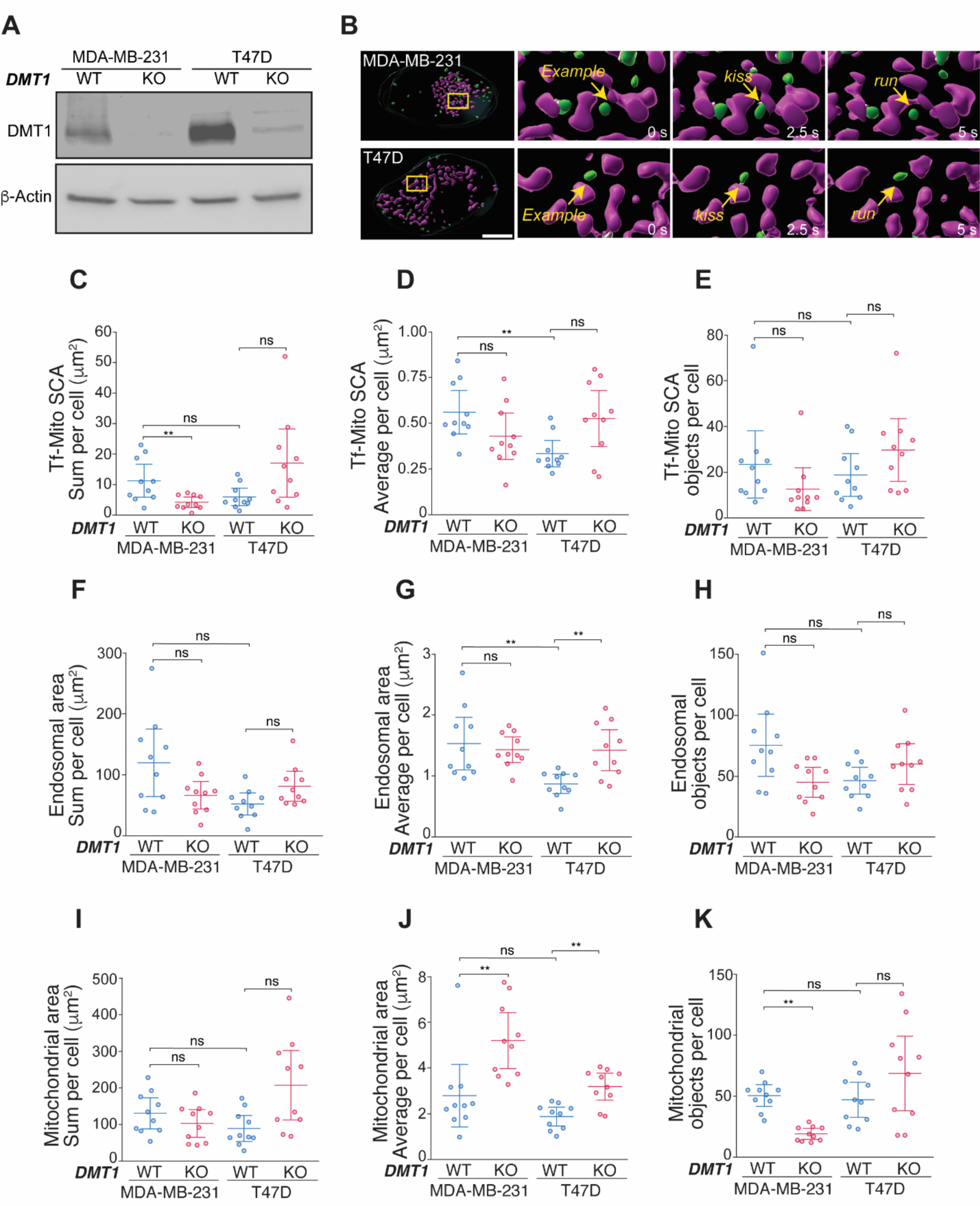
DMT1 knockout reduces endosome-mitochondria interactions in MDA-MB-231 but not in T47D cells. **(A)** CRISPR/Cas9 mediated DMT1 silencing was performed in these cell lines. DMT1 WT and KO cells were immunoblotted using anti-DMT1 antibody showing a significant decrease of DMT1 protein expression in both cell lines upon DMT1 silencing. **(B)** 3D reconstructions of Tf-containing endosomes (green), Mitotracker-labeled mitochondria (magenta) and endosome-mitochondria SCA (white) from live-cell imaging experiments (42.5 seconds videos; interval 2.5 seconds). Images were analyzed using IMARIS 9.6 software. Arrows indicate “kiss-and-run” endosome-mitochondria interactions. **(C)** A significant decrease of the total amount of Tf-Mito SCA sum per cell is observed upon DMT1 silencing in MDA-MB-231 but not in T47D. **(D-E)** SCA average **(D)** and number of Tf-Mito SCA objects **(E)** per cell are shown. **(F-K)** Total endosomal area sum **(F)**, endosomal area average **(G)**, number endosomal of objects **(H)**, total mitochondrial area sum **(I)**, mitochondrial area average **(J)** and mitochondrial objects **(K)** per cell are also shown. Parameters were calculated using data from 10 individual cells per condition, analyzed separately. One-way ANOVA with Bonferroni post-hoc test. **p<0.01. ns: non-significant (p>0.05). Scale bar = 10 μm.

Analyses of Tf object area measurements per cell showed no significant difference in the sum of endosomal area (**Figure 2F**), but a larger area on average of Tf objects in WT MDA-MB-231 compared to WT T47D cells was observed (**Figure 2G**); these results are in agreement with our previous published findings (Tubbesing et al., 2020). An increase in Tf endosomal area on average upon DMT1 ablation in T47D cells, but not in MDA-MB-231 cells, suggested a role for DMT1 in the regulation of endosomal morphology in T47D cells but not in MDA-MB-231 (**Figure 2G**). Mitochondrial 3D rendering shows that DMT1 silencing increased the average mitochondrial surface (**Figure 2J**) and reduced the number of mitochondrial objects in MDA-MB-231 cells (**Figure 2K**). These results suggest that DMT1 loss leads to increased mitochondrial tubulation and reduced fragmentation in these cells. In contrast, loss of DMT1 in T47D cells led to increased mitochondrial surface, but not to a reduced number of objects.

Lastly, although the number of SCA and endosomal objects per cell did not show differences between WT and DMT1 KO in both cell lines (**Figure 2E and 2H**), the number of mitochondrial objects was reduced upon DMT1 ablation in MDA-MB-231 but not in T47D (**Figure 2K**), thereby indicating that DMT1 appears to regulate both size and number of mitochondrial objects in MDA-MB-231 cells (**Figure 2J and 2K**). Overall, these data show that silencing of DMT1 decreases endosome-mitochondria interaction in MDA-MB-231 but not in T47D cells. In contrast, DMT1 ablation results in increased mitochondrial tubulation and decreased fragmentation in MDA-MB-231 cells. Given that DMT1 acts as a bridge between endosomes and mitochondria more frequently in MDA-MB-231 but not in T47D cells, MDA-MB-231 cells appear more sensitive to ablation of DMT1 for the establishment of endosome-mitochondria interactions.

### DMT1 modulates endosomal dynamics in MDA-MB-231 cells

In epithelial cells, we previously demonstrated that blocking iron release from Tf in the endosomal lumen can regulate both endosomal speed and duration of endosome-mitochondria transient interactions (Das et al., 2016). Here, we found that DMT1 silencing significantly decreases endosomal speed in MDA-MB-231, but not in T47D breast cancer cells (**Figure 3A and 3B**). Track length and track displacement length of endosomes were not affected by DMT1 silencing in both cell lines (**Figure 3C and 3D**). These results indicate that DMT1 regulates endosomal dynamics in MDA-MB-231 cells, which may be related to its role in modulating the levels of transient “kiss-and-run” endosome-mitochondria interactions (**Figure 2C**).

**Figure 3.**
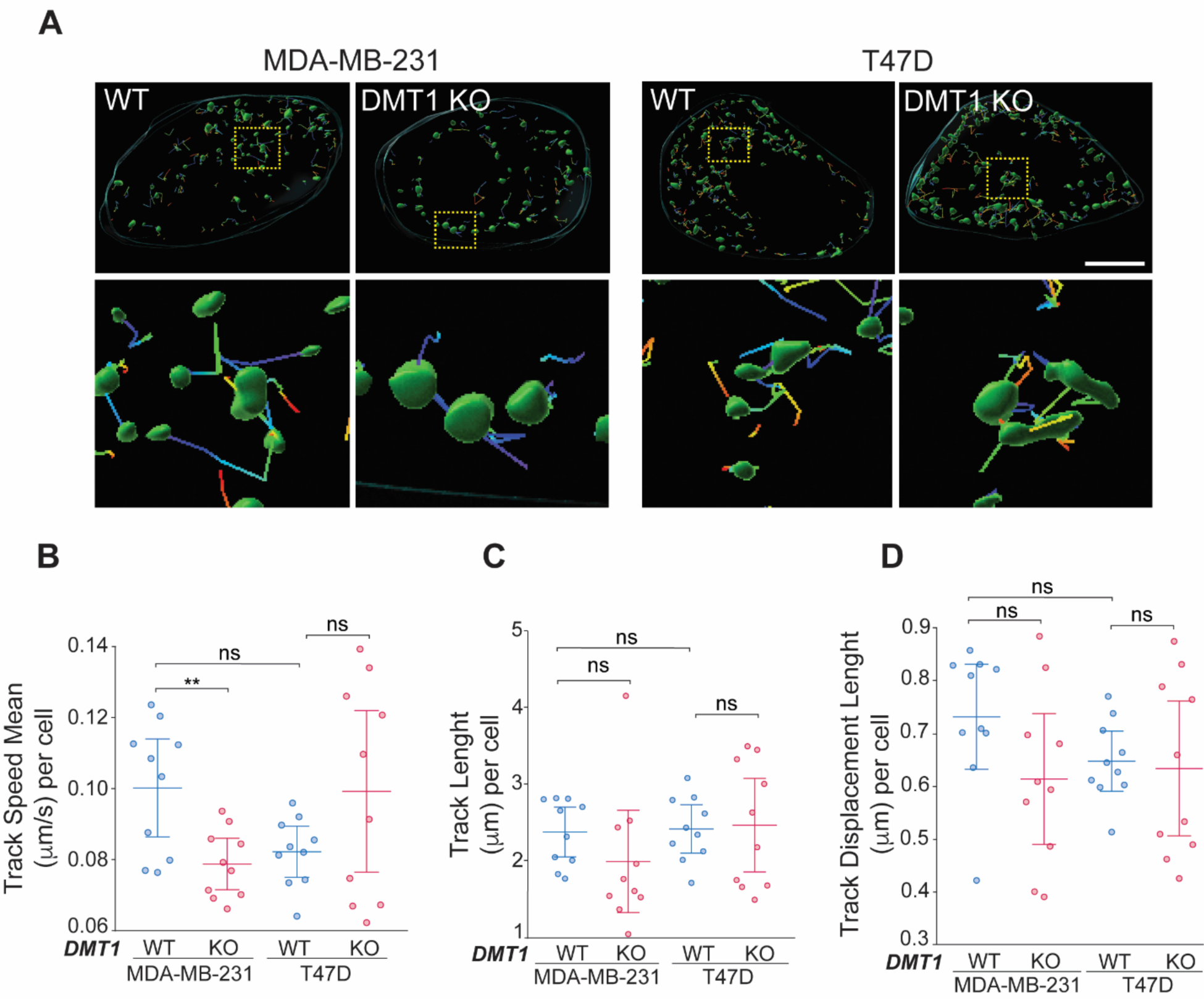
DMT1 silencing decreases endosomal speed in MDA-MB-231 but not in T47D cells. **(A)** Images shows endosomes and its endosomal tracks obtained from live cell imaging experiments. Yellow dotted squares indicate the corresponding magnified regions. Fluorescent images were 3D rendered and analyzed using IMARIS 9.6 software to quantify endosomal track speed mean **(B)**, track length **(C)** and track displacement length **(D)** per cell. One-way ANOVA with Bonferroni post-hoc test. **p<0.01. ns: non-significant (p>0.05). Scale bar = 10 μm.

### DMT1 regulates mitochondrial iron translocation in MDA-MB-231 cells

Previously we have shown that direct interactions between EE and mitochondria can facilitate mitochondrial iron translocation (MIT) in MDCK epithelial cells (Das et al., 2016). Here, we evaluated the involvement of DMT1 on MIT regulation in MDA-MB-231 and T47D breast cancer cells. We used the mitochondrial iron biosensor, RDA, to specifically measure MIT as described in (Das et al., 2016). RDA red-fluorescence undergoes rapid quenching upon iron translocation into the mitochondria (Rauen et al., 2007). Interestingly, we observed that DMT1 silencing in T47D cells did not significantly alter RDA fluorescence quenching compared to T47D WT parental cells (**Figure 4A and 4B**), indicating that DMT1 is not critical for MIT in T47D cells. However, DMT1 ablation in MDA-MB-231 led to a pronounced RDA quenching delay (**Figure 4C and 4D**), a phenotype that was partially rescued by re-expression of DMT1 in MDA-MB-231 DMT1 KO cells (**Figure 4C and 4D**). These results indicate that DMT1 regulates MIT in MDA-MB-231 cells but not in T47D cells. These experiments indicate a differential requirement of DMT1 for the maintenance of proper MIT levels in these two disparate breast cancer cell lines.

**Figure 4.**
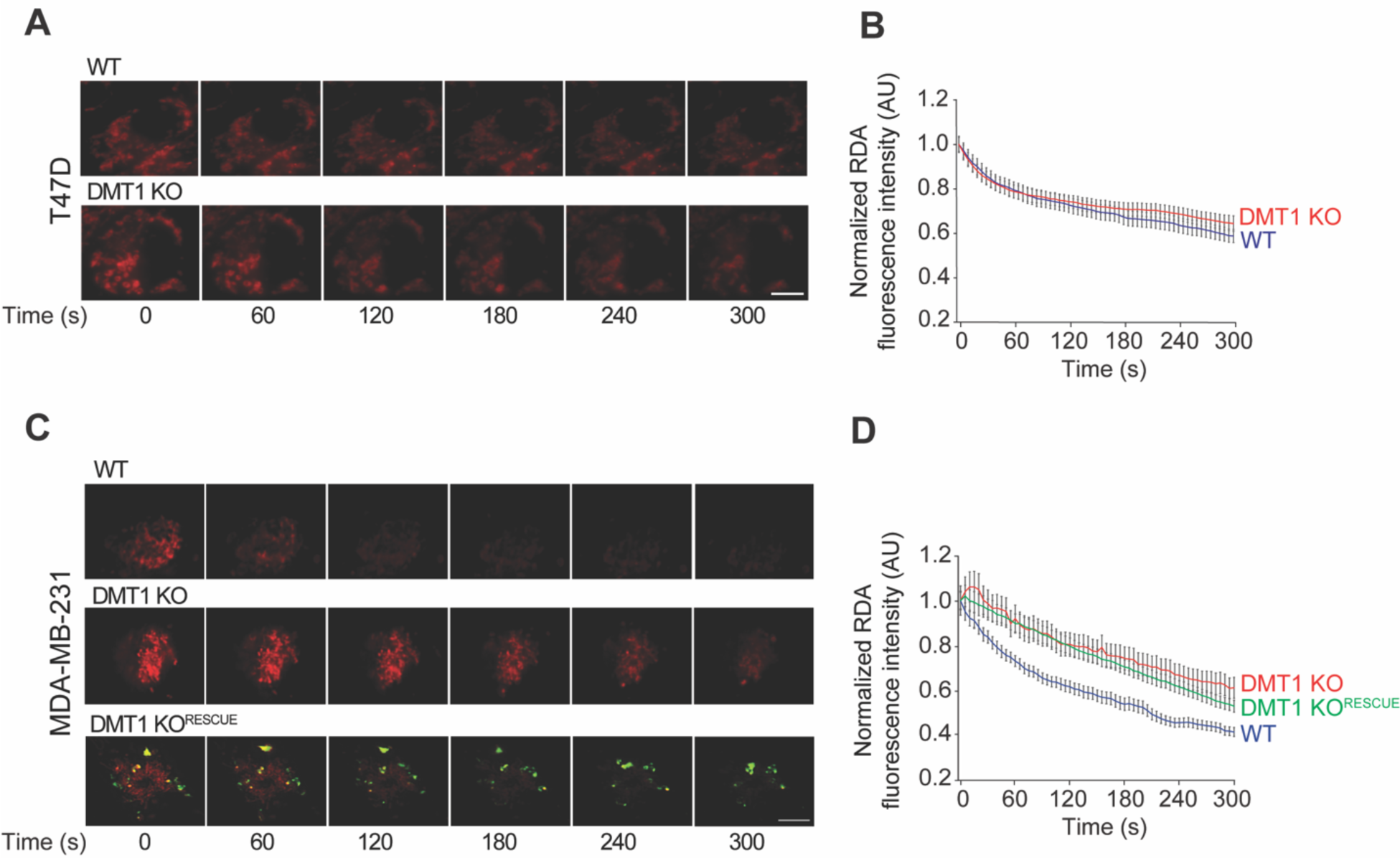
DMT1 regulates mitochondrial iron translocation (MIT) in MDA-MB-231 but not in T47D. Cells were incubated with RDA-MIT (a biosensor that localizes to mitochondria and undergoes rapid quenching upon iron binding) for 15 min and then subjected to live cell imaging. Fluorescent images and normalized fluorescence quantification graphs show RDA dequenching levels from 0 to 300 seconds (s). **(A-B)** DMT1 silencing (DMT1 KO) did not affect MIT in T47D. **(C-D)** In MDA-MB-231, DMT1 silencing (DMT1 KO) induces a delay in RDA dequenching indicating lower MIT. DMT1- GFP re-expression in DMT1 KO cells (DMT1 KO^Rescue^) partially rescued the decrease in MIT in DMT1 KO cells. RDA fluorescence intensity decay was analyzed using ImageJ in 10 cells per condition. Scale bar = 10 μm.

### DMT1 regulates the labile iron pool (LIP) in MDA-MB-231 breast cancer cells

As mentioned above, the main cellular regulators of iron import and export are TfR and FPN, respectively (Andrews and Schmidt, 2007). Here, we observed that DMT1 silencing decreased the expression of TfR and increased iron exporter FPN expression level in MDA-MB-231 cells (**Figure 5A and 5B**). To evaluate the effect of DMT1 silencing on LIP levels in MDA-MB-231 cells, we used the ferrous iron specific fluorescent dye FerroOrange, which reacts with LIP in cytoplasm. Interestingly, DMT1 ablation in MDA-MB-231 significantly increased LIP levels; WT LIP levels were rescued by DMT1 re-expression in DMT1 KO cells (**Figure 5C and 5D**). Mitochondrial iron chelation using mitochondrial targeted deferoxamine was shown to induce mitochondrial superoxide levels (Sandoval-Acuna et al., 2021). Mitochondria are the main source of ROS in the cells, and therefore tight regulation of ROS production and availability is critically controlled (Murphy, 2009). Here, we observed a significant increase in mitochondrial superoxide levels upon DMT1 silencing in MDA-MB-231 cells; again, re-expression of DMT1 in MDA-MB-231 DMT1 KO cells decreased mitochondrial superoxide to WT levels (**Figure S2B**). However, when mitochondrial membrane potential (MMP) was evaluated using the TMRM fluorescence probe, we observed a slight decrease in fluorescence levels, indicating a partial but not significant impairment in MMP upon DMT1 ablation (**Figure S2C**). Overall, these results indicate that DMT1 regulates LIP levels, TfR and FPN expression levels as well as the generation of superoxide by mitochondria in MDA-MB-231 cells.

**Figure 5.**
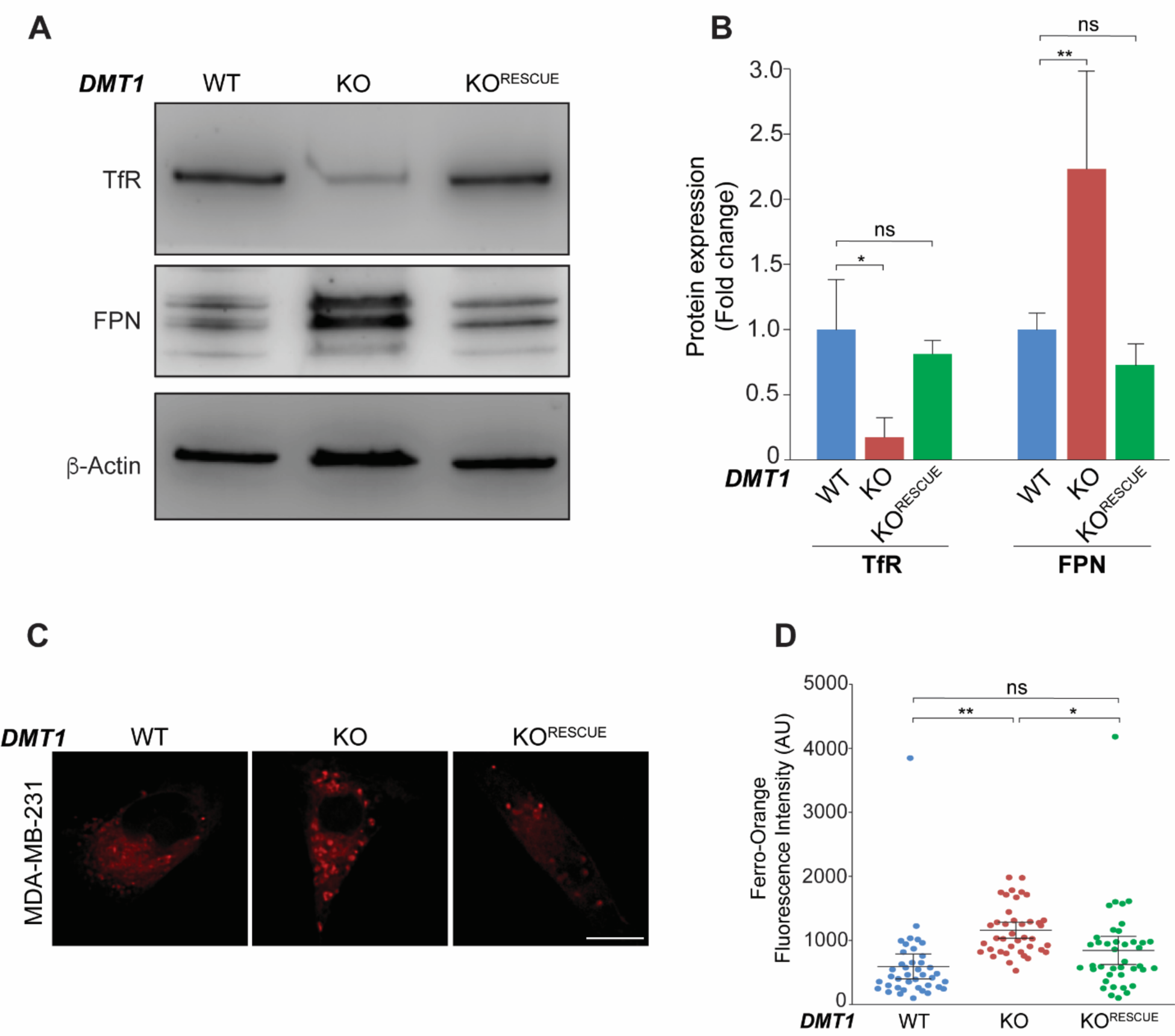
DMT1 regulates LIP levels and iron transport protein expression levels in MDA-MD-231 cells. **(A-B)** Immunoblot and normalized densitometry quantification showed levels of TfR and FPN in MDA-MB-231 WT, DMT1 KO and DMT1 KO^Rescue^ cells. β-Actin was used as a loading control. **(C-D)** Fluorescent images and normalized fluorescence quantification of Ferro-Orange dye in indicated cells. Graph shows quantification of fluorescence intensity in arbitrary units (AU). DMT1-GFP re-expression in DMT1 KO cells rescued the LIP increase induced by DMT1 silencing. Ferro-Orange fluorescence intensity was analyzed using ImageJ software in 40 cells per condition. One- way ANOVA with Bonferroni post-hoc test. *p<0.05; **p<0.01. ns: non-significant. Scale bar = 10 μm.

### DMT1 silencing induces PINK1/Parkin dependent mitophagy in MDA-MB-231 cells

Recent studies have demonstrated that iron chelation can induce mitophagy, which is the mitochondrial quality control process by which depolarized or damaged mitochondria are selectively subjected to degradation, with deleterious effects for cancer cells, including breast cancer (Bernardini et al., 2017; Hara et al., 2020; Palikaras et al., 2018; Sandoval- Acuna et al., 2021). Here, we evaluated the impact of DMT1 silencing on PINK1/Parkin mitophagy markers, mitochondrial ferritin (FTMT), as well as global autophagy marker LC3B in both T47D and MDA-MB-231 cells. Strikingly, in T47D none of these markers were altered upon DMT1 silencing (**Figure S1**) likely because in T47D DMT1 silencing does not affect MIT levels (**Figures 4A and 4B**). In contrast, DMT1 ablation in MDA-MB- 231 cells significantly induced PINK/Parkin as well as FTMT and LC3B expression levels (**Figures 6A and 6B**). Importantly, recovery to basal levels of all these markers was achieved by re-expression of DMT1 in MDA-MB-231 DMT1 KO cells (**Figures 6A and 6B**).

**Figure 6.**
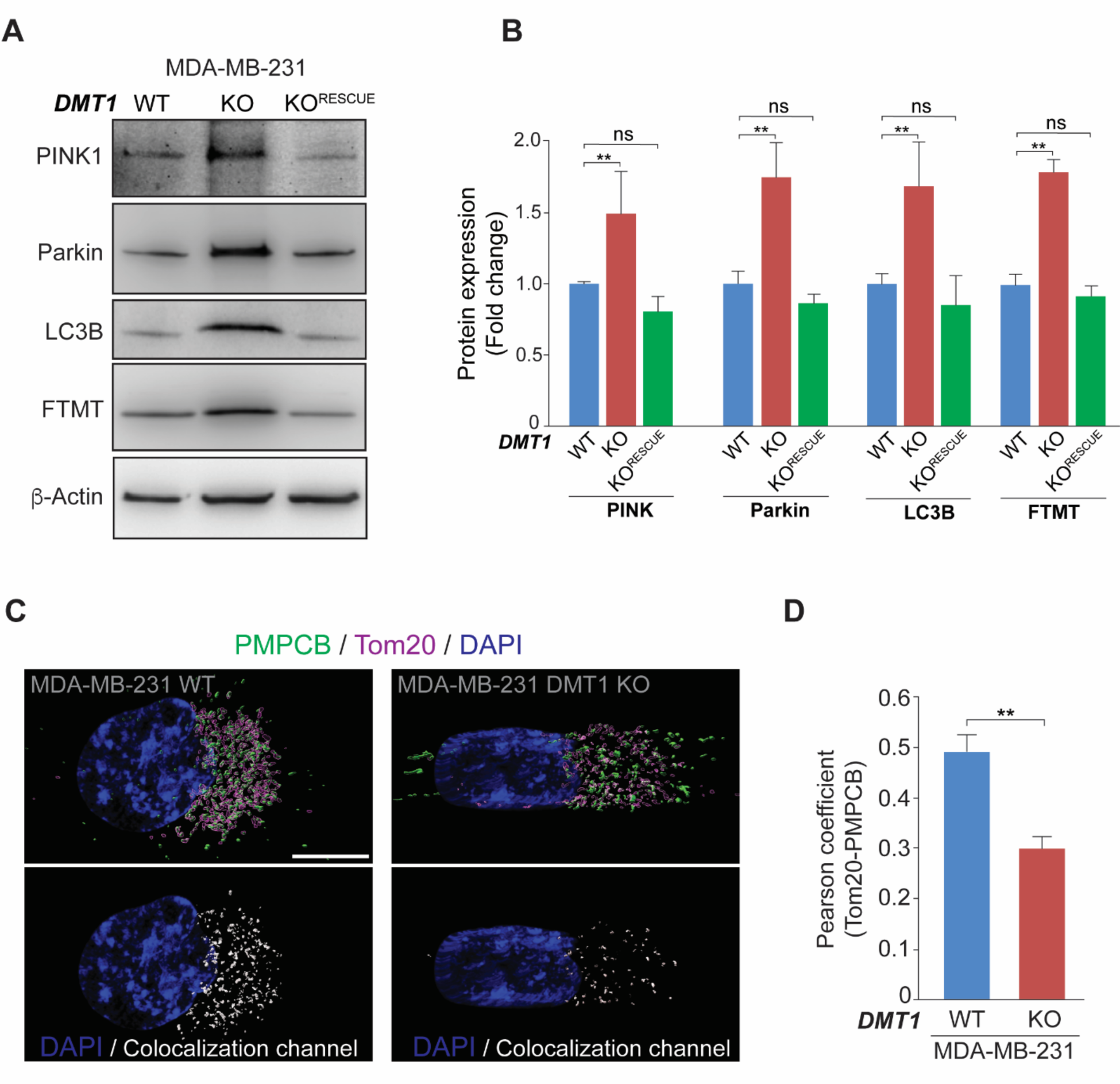
DMT1 silencing induces mitophagy and reduces PMPCB translocation to the OMM in MDA-MB-231 cells. **(A-B)** Immunoblots and normalized densitometry quantification of PINK1, Parkin, LC3B, and mitochondrial ferritin (FTMT) in MDA-MB-231 WT, DMT1 KO, and DMT1 KO^Rescue^ cells. ²-Actin was used as a loading control. **(C-D)** Immunofluorescence of Tom20 and PMPCB shows a decrease in colocalization between both proteins upon DMT1 silencing in MDA-MB-231. Pearson colocalization index was analyzed using IMARIS 9.6 software in 15 cells per condition. One-way ANOVA with Bonferroni post-hoc test (B) and unpaired *t*-test (D). *p<0.05; **p<0.01. ns: non-significant. Scale bar = 10 μm.

Interestingly, a recent study, where DMT1 (SLC11A2) was CRISPR-edited to endogenously express GFP, shows that DMT1 can interact with the peptidase PMPCB in HEK293 cells (Cho et al., 2022). PMPCB and PMPCA can establish the mitochondrial processing peptidase complex, where PMPCA regulates the import of proteins, such as PINK1, which can then be cleaved by PMPCB in the mitochondrial matrix (Quiros et al., 2015). PINK1 has been shown to be constitutively imported into the mitochondria, where it can be proteolytically degraded in a PMPCB-dependent manner (Greene et al., 2012). Failure to import or to degrade PINK1 can lead to its accumulation on the OMM, which leads to the hyperactivation of PINK1/Parkin-dependent mitophagy (Yang and Suda, 2018). Using immunofluorescence and Pearson coefficient colocalization analysis, we observed a significant decrease in the association between PMPCB and the OMM marker Tom20 upon DMT1 silencing in MDA-MB-231 cells (**Figures 6C and 6D**). These results indicate that DMT1 is involved in the crosstalk between PMPCB and OMM, and subsequent turnover of proteins, such as PINK1, thus, preventing mitophagy hyperactivation. Overall, our results reveal a differential dependence of DMT1 in MDA-MB-231 and T47D for PINK1/Parkin dependent mitophagy, which is likely connected to mitochondrial iron homeostasis in breast cancer cells.

### DMT1 regulates invasive migration and mitochondrial metabolism in MDA-MB-231 breast cancer cells

To investigate the association of DMT1 with breast cancer in patients, we evaluated DMT1 mRNA expression using data from clinically annotated patients’ genetic datasets of a normal breast tissue cohort [lnce (n = 21)] in comparison with cohorts of tumoral breast tissue samples [Clynes (n = 121) and Shaw (n = 44)]. We found a significant increase in DMT1 mRNA expression in tumoral breast tissue compared with normal breast tissue (**Figure S3A**). Moreover, by Kaplan Meier survival analysis of the breast invasive carcinoma patient TCGA genomic dataset, we found that higher expression of DMT1 is significantly associated with lower overall survival (**Figure S3B**). These genomic clinical data/analyses indicate an association between higher levels of DMT1 and breast cancer malignancy. RNA-sequencing was performed to analyze global gene transcription changes associated with DMT1 silencing in MDA-MB-231 WT versus DMT1 KO cells. Our RNA sequencing data revealed a total of 411 differentially expressed genes upon DMT1 silencing in MDA-MB-231 cells (131 upregulated and 280 downregulated), when we used the cutoff ± 1.5 log2FoldChange in the DEG analysis (**Figure 7A**, **Table S3**). Gene ontology (GO) biological processes evaluation showed that DMT1 silencing affected important cancer-related processes such as cell adhesion, cell migration, and cell motility (**Figure 7B**). Additionally using the GO database associated to diseases (Diseases GLAD4) we observed that DMT1 silencing in MDA-MB-231 decreases significantly GO terms “carcinoma” and “epithelial cancers” (**Figures S4A, S4B and S4C**). Furthermore, integrated system for motif activity response analysis (ISMARA), that evaluates computationally predicted regulatory sites for transcription factors, revealed NRF1 transcription factor activity as the top increase upon DMT1 silencing in MDA-MB-231 cells (**Figure S4D**). Interestingly, NRF1 may contribute to therapeutic strategies in cancer by reactivating immunity and sensitizing tumoral cells to apoptosis (Ma et al., 2019). In contrast, the top decrease in transcription factor activity upon DMT1 KO in MDA-MB-231 was HIF2A (**Figure S4D**), which has been associated with enhanced breast tumoral cells dissemination (Todd et al., 2021). Altogether, these data prompted us to evaluate invasive migration in 3D matrices, an important feature of aggressive cancer cells that is essential for metastasis initiation. Inverted invasion assays in Matrigel plugs supplemented with fibronectin showed a significant decrease of invasive migration upon DMT1 silencing in MDA-MB-231 cells that was partially recovered upon DMT1 re-expression in DMT1 KO cells (**Figure 7C and 7D**).

**Figure 7.**
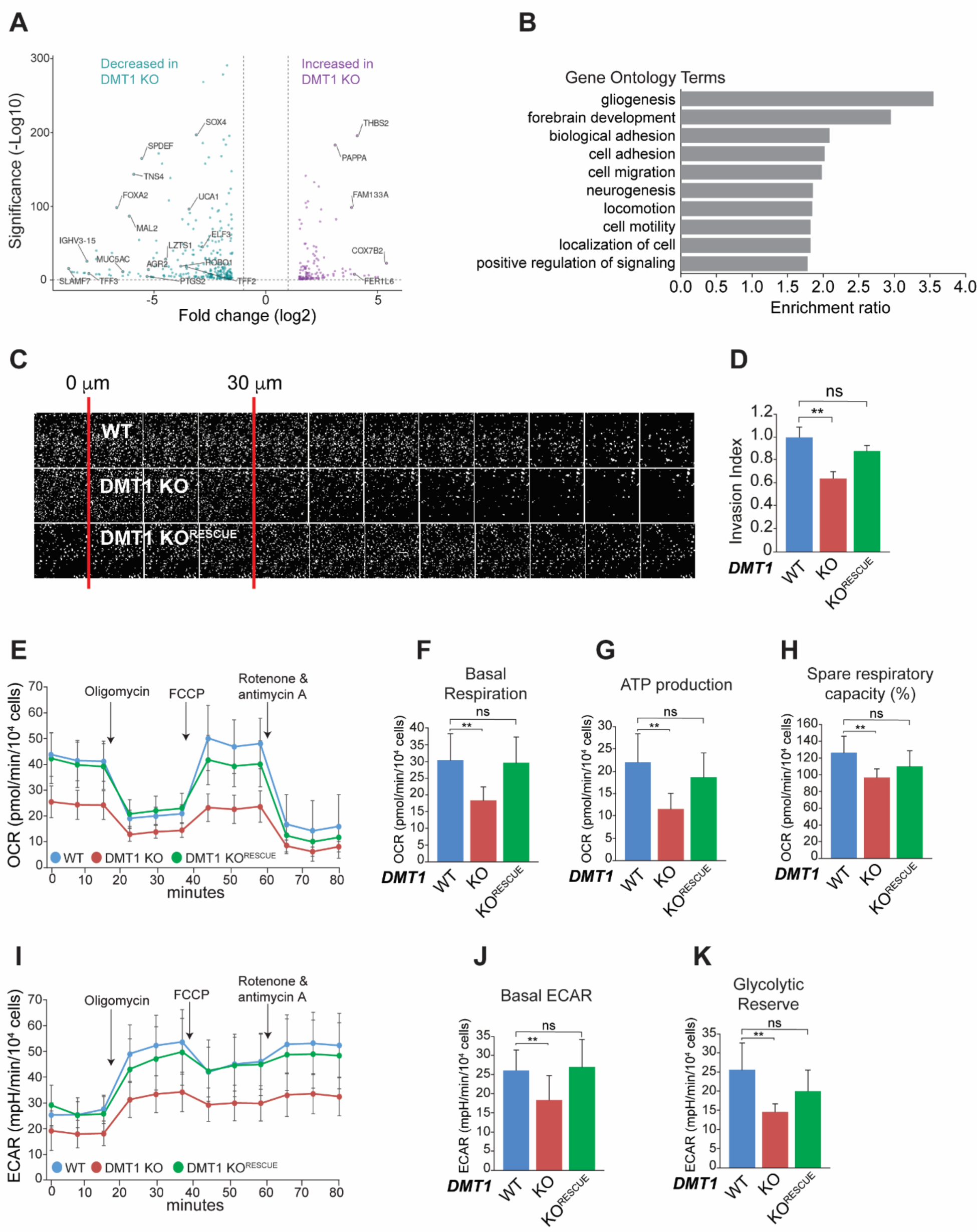
DMT1 regulates mitochondrial bioenergetics and invasive migration in MDA-MB-231. **(A)** RNA sequencing analyses of MDA-MB-231 WT and DMT1 KO. Volcano Plot shows downregulated (green) and upregulated (purple) genes upon DMT1 silencing. **(B)** Biological processes gene ontology (GO) analysis. **(C-D)** Inverted invasion assay shows a significant decrease in DMT1 KO cells’ invasive migration, phenotype that is rescued by re-expression of DMT1. **(E)** Oxygen consumption rate (OCR) and **(I)** extracellular acidification rate (ECAR) in response to sequential treatment with oligomycin, FCCP and Rotenone & Antimycin A of MDA-MB-231 WT, DMT1 KO and DMT1 KO^RESCUE^ cells are shown. Data was normalized for number of the cells (40,000 cells/well) that were stained using Hoechst at the end of the assay (n=3, 6-10 replicates per condition). Basal respiration **(F)**, ATP production **(G),** and spare respiratory capacity **(H)** were calculated using OCR data. Basal ECAR **(J)** and glycolytic reserve **(K)** were calculated using ECAR data. Bar charts: one-way ANOVA with Bonferroni post-hoc test. *p<0.05; **p<0.01. ns: non-significant. Scale bar = 100 μm.

**Figure 8.**
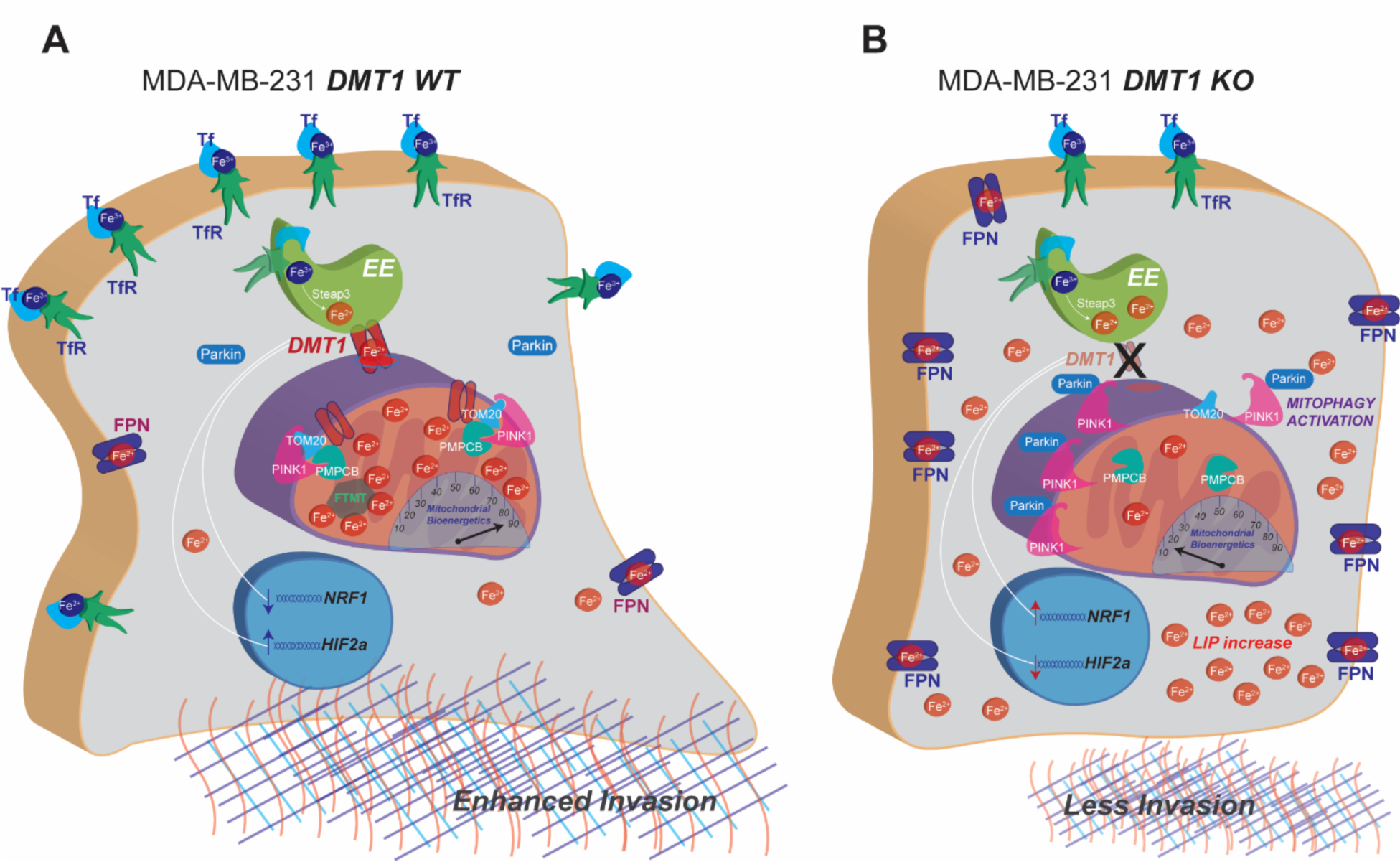
Schematic representation of DMT1 silencing effects in MDA-MB-231 cells. **(A)** In MDA-MB-231 WT cells, ferric iron (Fe^3+^) binds to Tf and iron-bound Tf-TfR complexes are taken into cells. Fe^3+^ is released into the lumen of early endosomes (EE) where is reduced by Steap3 to its Fe^2+^ form. Then DMT1 can translocate Fe^2+^ to mitochondria, where iron is accumulated. In mitochondria, Fe^2+^ is either stored in mitochondrial ferritin (FTMT) or used for mitochondrial bioenergetics, typical of invasive cancer cells. PMPCB, a DMT1 interactor, associates with TOM20 at the OMM, preventing PINK accumulation. **(B)** Upon DMT1 silencing, mitochondrial translocation of Fe^2+^ is reduced, as are the mitochondrial bioenergetics and invasive capacity; Fe^2+^ accumulates in the cytoplasm increasing LIP levels. TfR levels are decreased and FPN levels are increased, likely as a compensatory mechanism of rising LIP levels. PMPCB dissociates from TOM20, preventing PINK1 turnover and activating Parkin recruitment and mitophagy. Transcription factors *NRF1* and *HIF2A* are activated and inhibited, respectively.

Invasive cancer cells can notably enhance and adapt their energy metabolism to support invasion, a highly-demanding energy process (Garde and Sherwood, 2021). For example, metastatic breast cancer cells cultured in extracellular matrix (ECM) can increase glucose uptake and ATP production (Zanotelli et al., 2018). Mitochondria can also localize towards energy-demanding leading edge and protrusion in several cancer cell types with the capacity to adapt ATP metabolism thus overcoming ECM barriers (Garde and Sherwood, 2021). To determine the role of DMT1 in regulating MDA-MB-231 cell bioenergetics, we measured oxygen consumption (OCR) and extracellular acidification rates (ECAR) in MDA-MB-231 with or without loss of DMT1. Cells were sequentially treated with oligomycin, FCCP, and rotenone and antimycin A to interrogate several aspects of mitochondrial respiration and their impact on aerobic glycolysis. We found that loss of DMT1 was associated with a decrease in both mitochondrial respiration (decrease basal OCR) and aerobic glycolysis (decrease basal ECAR) upon loss of DMT1 (**Figures 7E and 7I**). Both basal OCR and ECAR were restored to levels of parental WT cells following reintroduction of DMT1 into MDA-MB-231 DMT1 KO cells (**Figures 7E, 7I, 7F, and 7J**). ATP production (**Figure 7G**), spare respiratory capacity (**Figure 7H**), basal ECAR (**Figure 7J**) and glycolytic reserve (**Figure 7K**) decreased upon DMT1 silencing, and returned to basal levels upon DMT1 rescue in KO cells. Thus, we conclude that in MDA-MB-231 breast cancer cells, DMT1 is necessary for invasive migration capacity and proper mitochondrial respiration and glycolysis.

## Discussion

Collectively, our present work demonstrates that DMT1 plays a key role in intracellular and mitochondrial iron homeostasis as well as mitophagy, mitochondrial bioenergetics and invasive capacity, by regulating endosome-mitochondria interactions in invasive triple negative (MDA-MB-231) but not in non-invasive luminal A (T47D) breast cancer cells. We observed that DMT1 silencing induces the deregulation of iron homeostasis via reduction of endosome-mitochondria interactions, MIT blockage, and higher LIP levels. Concomitantly, DMT1 ablation results in both a compensatory phenotype of increase- export/decrease-import of iron and the induction of PINK1/Parkin-dependent mitophagy in invasive MDA-MB-231, but not in non-invasive T47D breast cancer cells. Furthermore, our analyses from publicly available genomic databases revealed a significant association of DMT1 with malignancy in invasive breast carcinoma patients. Concordantly, RNA sequencing data from MDA-MB-231 WT versus DMT1 KO cells corroborated DMT1 as an invasive breast cancer driver gene. Moreover, invasion in 3D matrices and mitochondrial respiration, and aerobic glycolysis were severely impaired upon DMT1 silencing in MDA-MB-231 cells and recovered by DMT1 rescue in DMT1 KO cells. Therefore, we propose that DMT1 acts as a crucial regulatory factor in invasive breast carcinoma.

T47D and MDA-MB-231 breast cancer cells are representative of two different types of breast cancer. MDA-MB-231 is a well characterized cellular model of triple-negative breast cancer, showing a mesenchymal phenotype, which is consistent with their capacity to invade *in vitro* and to generate spontaneous metastases when orthotopically implanted in immunodeficient mice (Holliday and Speirs, 2011). Conversely, the luminal A breast cancer cell line T47D, forms highly cell-cell adhesion cohesive clusters, representative of a more epithelial non-invasive phenotype. Moreover, T47D’s ability to form large and aggressive tumor xenografts depends on the addition of estrogen (Holliday and Speirs, 2011). Our previous work demonstrated a remarkable difference between these two cell lines in terms of iron-related protein expression, endosomal trafficking, transferrin recycling, and endosomal iron retention/release properties (Khoo et al., 2020; Tubbesing et al., 2020). Here we showed a differential DMT1 requirement for both endosome-mitochondria interactions and MIT when MDA-MB-231 and T47D breast cancer cells were compared. Specifically, DMT1 regulates endosome-mitochondria “kiss-and-run” transient interactions as well as endosomal dynamics and MIT levels in MDA-MB-231 but not in T47D cells. Triple immunofluorescence of endosomes, DMT1 and mitochondria showed that DMT1 frequently bridges endosomes and mitochondria in MDA-MB-231 but not in T47D cells. These findings indicate that invasive MDA-MB-231 cells are dependent on DMT1 for the proper management of MIT and its associated metabolism. In contrast, DMT1 is not required for the regulation of these processes in T47D, indicating the existence of multiple iron regulatory pathways in breast cancer cells. Even though MDA-MB-231 cells are more aggressive than T47D, they may be particularly vulnerable to DMT1-dependent mitochondrial iron metabolism disruption.

Our previous work showed that endosomes are larger and more peri-nuclear in MDA-MB- 231 than in T47D (Tubbesing et al., 2020). Interestingly, mitochondria also tend to be concentrated in the peri-nuclear region in MDA-MB-231 cells. Moreover, MDA-MB-231 cells show a delay in iron release from Tf in endosomes as compared to T47D (Khoo et al., 2020). These previous results agree with our current findings that show a specific vulnerability of MDA-MB-231 in the DMT1-dependent mitochondrial iron management, when compared to less aggressive T47D cells. The notable differences in endosomal characteristics between these two breast cancer cell types can be explained by numerous variables such as genetic background, level of differentiation, differential activity of enzymes and metabolic pathways, or presence of mutations in signaling pathways such as PI3K that are important for endocytic trafficking and regulation of endosomal maturation (Wallroth and Haucke, 2018). Nevertheless, an interesting hypothesis is that the different intracellular distribution of endosomes and mitochondria within the two breast cancer cell lines analyzed in this study, can be the basis of a novel invasive cancer cell’s vulnerability related to iron homeostasis. A caveat of our work is that, although we compared cell lines from two different breast cancer subtypes, we used only two cell lines, which prevents us from a generalization of these mechanisms in breast cancer cells. Ongoing and future work in our lab comparing organelle distribution and its features as well as the evaluation of other candidates for MIT regulation will advance our understanding of the regulation of this specific targeting of mitochondrial iron metabolism in aggressive breast cancer cells.

PINK1/Parkin mediated mitophagy can regulate breast cancer malignancy (Bernardini et al., 2017). Interestingly, iron chelation specifically increases the expression of mitochondrial ferritin (FTMT) and the translocation of its precursor to the OMM, inducing mitophagy and suppressing tumor development (Hara et al., 2020). Moreover, it has been recently described that mitochondrial iron chelation activates mitophagy, impairing mitochondrial metabolism and decreasing invasion and tumoral growth both *in vitro* and *in vivo* (Sandoval-Acuna et al., 2021). Specifically, a specific mitochondrial iron chelation, using a novel mitochondrial targeted deferoxamine (mitoDFO), induces PINK1- dependent mitophagy in both non-invasive MCF7 and invasive MDA-MB-231 breast cancer cells (Sandoval-Acuna et al., 2021). Notably, here we found that DMT1 silencing induces PINK1/Parkin-dependent mitophagy in invasive MDA-MB-231 but not in non-invasive T47D breast cancer cells. Interestingly, Sandoval-Acuña et al., showed only slight differences in TfR and FPN expression upon mitoDFO treatment, suggesting that LIP in cytosol is not significantly affected by mitoDFO. In our work, firstly we showed that upon DMT1 silencing in MDA-MB-231 cells TfR is downregulated, FPN is upregulated, and cytosolic LIP is increased. Secondly, DMT1 silencing in MDA-MB-231 cells also increased mitochondria superoxide production, although without affecting mitochondrial membrane potential. Thirdly, we found that DMT1 is required for the appropriate colocalization between Tom20 and the PINK1 turnover regulator PMPCB (Yang and Suda, 2018), a recently described DMT1 interactor (Cho et al., 2022). Silencing of DMT1 in MDA-MB-231 shifted the oncogenic high-TfR/low-FPN phenotype to the low-TfR/high- FPN breast tumor suppressor phenotype (Miller et al., 2011). The combination of the increasing LIP and ROS levels via MIT blockage upon DMT1 silencing, with the experiments demonstrating the role of DMT1 in regulating mitochondrial respiration and aerobic glycolysis, could explain the significant decrease on invasive migration upon DMT1 ablation in MDA-MB-231 breast cancer cells. Of note, these negative effects in mitochondrial bioenergetics and invasive capacity are achieved by a highly specific and targeted disruption of DMT1-dependent iron metabolism.

Expression of NRF1 (nuclear factor erythroid 2-like 1), a member of the cap‘n’collar basic region-leucine zipper transcription factor family involved in regulation of oxidative stress, is reduced in breast cancer (Sekine and Motohashi, 2021). Moreover, NRF1 blocking may play a significant role in tumor suppression, contributing to therapeutic strategies in cancer via immune reactivation and tumoral cells sensitizing to apoptosis (Ma et al., 2019; Sekine and Motohashi, 2021). Interestingly, our RNA sequencing data from MDA-MB- 231 WT versus DMT1 KO cells shows a significant increase in transcriptional activity of NRF1 target genes. In contrast, the top decrease in transcription factor activity was the gene HIF2A, which has been associated with poor outcome in invasive breast carcinoma (Helczynska et al., 2008). Together with the strong association of DMT1 expression with breast cancer, as well as with a decrease in overall survival in invasive breast carcinomas patients, our results highlight DMT1 as an important driver in this aggressive type of cancer.

In summary, this study demonstrates a critical requirement of DMT1 for an adequate mitochondrial metabolism as well as invasive capacity, due to DMT1’s role in endosome-mitochondria transient interactions, MIT, mitophagy, and cellular iron metabolism/transport regulation, specifically in invasive breast cancer cells. Future experiments seek to corroborate the hypothesis of a specific role of DMT1 as a particular and highly specific Achilles’ heel for invasive breast cancer cells that could be exploited therapeutically.

## Materials and Methods

### Cell culture

MDA-MB-231, T47D and HEK293T cell lines were obtained from ATCC (Manassas, VA, USA) and grown at 37°C in a humidified atmosphere containing 5%-CO_2_. Cells were cultured in DMEM (ThermoFisher) supplemented with 10% fetal bovine serum (FBS) (ATCC), 100 units/ml penicillin, 100 ug/ml streptomycin and 4 mM L-glutamine. Cells were used at passage number equal or lower than 10, and tested routinely for mycoplasma contamination using PCR.

### CRISPR/Cas9 mediated DMT1 silencing

Silencing of DMT1 in MDA-MB-231 and T47D cell lines was performed using lentiCRISPRv2 plasmid containing the DMT1 (SLC11A2) targeting sgRNA sequence 5’-TGAGAAGATCTCCATTCCTG-3’. Lentiviral particles were produced in HEK293T cells by transfection with 0.2 µg of pCMV-VSV-G (Addgene, #8454, MA, USA), 2 µg of psPAX2 (Addgene, #12260, MA, USA) and 2 µg of lentiCRISPRv2 sgRNA DMT1. After 48 h, the supernatant of transfected cells was collected and filtered using a 0.45-µm PES filter. Lentiviral suspension was immediately used for transduction or stored at −80°C. A 1:1 lentiviral vector suspension containing 10 µg of polybrene (SCBT) was added drop by drop to either MDA-MB-231 or T47D cells and incubated for 72 hours. For antibiotic selection, 1 µg/mL of puromycin (Gibco, USA) was used. The cells were cultured until all the non-transduced control cells died (2-3 days) plus 2 more days in antibiotic selection media. For rescue of DMT1 expression in MDA-MB-231 DMT1 KO, cells were transfected with plasmid pCIneo-DMT1-GFP (N-terminal GFP, DMT1 isoform 2, kindly provided by Dr. Jerry Kaplan, Department of Pathology, University of Utah Health Sciences Center, Salt Lake City, UT) and cells were further selected with Neomycin (100 μg/ml) until all non-treated control cells died (2 weeks) plus 2 more days in antibiotic selection media.

### Indirect immunofluorescence

Immunofluorescence experiments were performed as previously described (Tubbesing et al., 2020). Briefly, cells were seeded into Ibidi 8 well μ-plates (Ibidi, #80827) and grown overnight at 37°C in a 5% CO_2_ environment. Cells were fixed for 15 min with 4% paraformaldehyde (PFA) at 37°C, permeabilized with 0.2% TritonX-100 in PBS for 15 min at room temperature, blocked for 90 min on a gentle rocker-shaker in blocking buffer (1% bovine serum albumin (BSA), 0.5% fish skin gelatin (FSG), 0.2% Triton X-100), and immunostained with primary antibodies (Supplementary table S1) overnight at 4°C, and then incubated with secondary antibodies for 1 h at room temperature, post-fixated with 4% PFA for 10 min, and stained with DAPI (1μg/ml) for 15 min. All solutions were 0.45 μm syringe filtered. Images were acquired using Thunder microscope (Leica) with oil immersion 63X/NA 1.4 objective, subsequently subjected to integrated computational clearing deconvolution process (LASX software) and then 3D-rendered and further analyzed using IMARIS 9.6 software (Bitplane). Colocalization Pearson coefficient was calculated using IMARIS integrated “coloc” tool (Tubbesing et al., 2020).

### Live cell imaging

For live cell imaging, cells were seeded in N° 1.5 coverslips-bottom 35-mm Petri dishes coated with poly-D-lysine (MatTek Corporation). Imaging medium comprised phenol red– free and serum-free DMEM supplemented with 4 mM L-glutamine. Live mitochondrial and early endosomes (EE) staining was achieved with 100 nM MitoTracker Red (Molecular Probes) and 2-min pulse of 50 µg/ml Tf AF488 (Invitrogen), respectively. Before imaging, the cells were briefly washed and chased with imaging media for another 2 min. Starting at time point 5–6 min after Tf pulse, time-lapse videos of 45-s duration were acquired. Assay was performed under CO_2_ and temperature-control within the live-cell incubation chamber of the Thunder microscope (Leica). Mitochondrial iron translocation (MIT) was measured incubating live cells with 100 nM iron sensor dye RDA (a gift from Dr. Ursula Rauen, Institute of Physiological Chemistry, University Hospital Essen, Essen, Germany) for 15 min at 37°C in cell incubator and then subjected to live cell imaging for 5 min at time interval of 5-s performed under CO_2_ and temperature-controlled incubation chamber of the Thunder microscope using oil immersion 63X/NA 1.4 objective (Leica). Labile iron pool, mitochondrial superoxide levels and mitochondrial membrane potential were evaluated by fluorescence imaging using FerroOrange (Dojindo), MitoSOX Red (Thermo Fisher Scientific) and tetramethylrhodamine ester TMRM (Thermo Fisher Scientific), respectively. Briefly, live cells were plated as described above and incubated at 37°C with either 1μM FerroOrange, 1μM MitoSOX or 100nM TMRM 100 for 30 min, respectively. Live cells were imaged using the Thunder microscope with oil immersion 63X/NA 1.4 objective (Leica). Fluorescence dequenching of RDA and mean intensity of FerroOrange, MitoSOX red or TMRM was analyzed in 40 cells per condition using ImageJ software (NIH).

### 3D rendering and surface contact area (SCA) analyses

Images from indirect immunofluorescence and live cell imaging experiments were 3D rendered using IMARIS 9.6 (Bitplane). Briefly, the surface of mitochondria was determined according to either the Mitotracker or anti-Tom20 immunostaining, while endosomal structures were obtained from Tf uptake data. DMT1 surface was obtained from anti-DMT1 immunostaining. For SCA measurements between organelle markers, (https://imaris.oxinst.com/open/view/surface-surface-contact-area) was used. The morphology properties of mitochondria, Tf, DMT1 and SCA properties were measured using IMARIS 9.6 surface module. Data from 10 rendered cells for each experimental condition was analyzed.

### Immunoblotting

Cells were lysed in immunoblotting buffer (25 mM HEPES, 150 mM NaCl, 1 mM MgCl2, 0.4% NP-40), supplemented with anti-protease and anti-phosphatase mixture (Millipore) at 4°C. Protein lysates were clarified by centrifugation. Proteins lysates for western blotting were separated by SDS-PAGE, transferred to PVDF membranes (Biorad), and then subjected to standard immunoblotting procedures as previously described (Tubbesing et al., 2020).

### Cellular bioenergetics

Cells were seeded (2 × 10^4^ cells/well) in XF96-well microplates (Agilent Technologies) and cultured at 37°C 5% CO_2_ for 24 h. The day before the assay, XF sensor cartridges were hydrated. One hour before the assay, culture media was replaced with DMEM XF base medium pH 7.4 supplemented with glutamine and glucose (Sigma-Aldrich). Oxygen consumption rate (OCR) and extracellular acidification rate (ECAR) were measured at 37°C on an XF Extracellular Flux Analyzer (Agilent Technologies) using the Cell MitoStress Test Kit (Seahorse Bioscience), according to manufacturer instructions. The baseline (basal) OCR was measured three times before, and then three times after each sequential injection of oligomycin (1 μM), carbonyl cyanide 4-(trifluoromethoxy) phenylhydrazone (FCCP) (0.5 μM), and rotenone and antimycin A (both 0.5 μM) and this last injection port was used also for nuclei stain using Hoechst. Upon completion of the assay, the microplate was imaged using Cytation5 microplate reader (BioTek) for nuclei stain detection. Assay was normalized by number of cells in each well. Data generated by the XF Extracellular Flux Analyzer (Agilent Technologies) were exported and analyzed using Microsoft Excel.

### Invasion assay

Invasive migration capacity was measured using an inverted invasion assay, as previously described (Barra et al., 2021; Hennigan et al., 1994). Briefly, Matrigel (Corning) 5 mg/mL, mixed with fibronectin (25 μg/mL), was polymerized within 12-well Transwell™ polycarbonate filter inserts (8-μm pore size; Corning) for 1 h at 37°C. Inserts were inverted, and 8 × 10^4^ cells were seeded at filter bottom and allowed to attach for 3–6 h at 37°C in 5% CO_2_ and then returned to the right side up. Serum free media was added to the wells of the Transwell plate, and media with FBS 10% plus 20 ng/mL EGF was added on top of the Matrigel plug, and incubated at 37°C in 5% CO_2_. After 5 days of incubation, Matrigel plugs were fixed with 4% PFA, treated with 0.2% Triton X-100 and stained with DAPI. Cells at the bottom of the filter that failed to migrate through the filter were removed using tissue paper. Images at 10-μm intervals were taken with Thunder microscope (Leica) using a 16X objective. ImageJ software (NIH) was used to determine the integrated density of each image section to calculate the invasion index = (∑ integrated density of first 30 µM)/(∑ integrated density of invasion) that was expressed as a fold change.

### RNA sequencing and genomic data analysis

Total RNA was isolated in triplicate form MDA-MB-231 WT and MDA-MB-231 DMT1 KO using TRIzol according to manufacturer instructions. Quality control of RNA determines that RIN number of all 6 samples processed was >8.6. RNA library preparation was performed using a polyA selection method. RNA sequencing was performed using Illumina HiSeq system in a 2 x 150-bp configuration (single index, per lane) by GENEWIZ. Starting from the raw .fastq files, the reads were mapped against the human reference genome (hg19Full) using the RNA STAR aligner (Dobin et al., 2013). Read counting was performed with htseqcount (Anders et al., 2015). Differential gene expression was analyzed using DESeq2 (Love et al., 2014). Significance was defined as P value<0.05 after adjustment for multiple testing. FastQC quality check analysis of the MDA-MB-231 WT and DMT1 KO samples in triplicate, achieved a high-quality score. Principal component analysis (PCA) indicates a distinct gene expression profile between conditions analyzed. PC1 accounts for 93.74% of transcriptional variance resulting from DMT1 silencing in MDA-MB-231 cells. PC2 accounts for 5.38% of differences into both groups, most likely due intrinsic cell heterogeneity. Other components of this analysis account for less than 1% of variance. Functional enrichment analysis was performed using WebGestalt (WEB-based Gene Set Analysis Tollkit, http://www.webgestalt.org) (Wang et al., 2017). ISMARA (Integrated System for Motif Activity Response Analysis) (Balwierz et al., 2014) was employed to estimate the key transcription factors associated with the observed differential gene expression of our RNA sequencing data from MDA-MB-231 WT and MDA-MB-231 DMT1 KO cells.

### Statistical Analyses

One-way ANOVA with Bonferroni post-hoc test, or unpaired two-tailed Student t-tests were used for experimental data analysis. Data are presented as mean with 95% confidence interval or SD were indicated. All experiments were performed with a minimum of three independent replicates. Number of cells analyzed for imaging analyses are indicated in respective figure legends. *p*-values less than 0.05 were considered statistically significant (*p < 0.05; **p < 0.01). Data were analyzed using GraphPad Prism 6.0 or Microsoft Excel and data visualization were generated using PlotsOfData web app (Postma and Goedhart, 2019) and Adobe Illustrator (Adobe).

## Acknowledgements

The authors thank Dr. Jerry Kaplan (Department of Pathology, University of Utah Health Sciences Center, Salt Lake City, UT) for providing DMT1-GFP plasmid and Dr. Ursula Rauen (Institute of Physiological Chemistry, University Hospital Essen, Essen, Germany) for the mitochondrial iron biosensor RDA. We also thank the members of Barroso, Jourd’heuil, Adam, Lamar labs at AMC, and the Corr and Intes labs at RPI for their help and support. We acknowledge funding support from National Institute of Health (NIH) grants R01 CA233188 and R01 CA207725 (to MB), NIH RO1HL142807-01 (to DJ).

## Author Contributions

J Barra: investigation, formal analysis, methodology, and writing-original draft, review, and editing.

I Crosbourne: investigation and formal analysis.

L Wang: investigation and formal analysis.

R Bossardi-Ramos: investigation and formal analysis.

F Jourd‘heuil: investigation and formal analysis.

I Nelson: investigation and formal analysis.

AP Adam: formal analysis and writing-original draft.

DT Corr: formal analysis and writing-original draft.

D Jourd‘heuil: formal analysis and writing-original draft.

M Barroso: conceptualization, investigation, data curation, supervising, funding acquisition, formal analysis, writing-original draft, review, and editing.

---

**Figure Supplementary S1.**
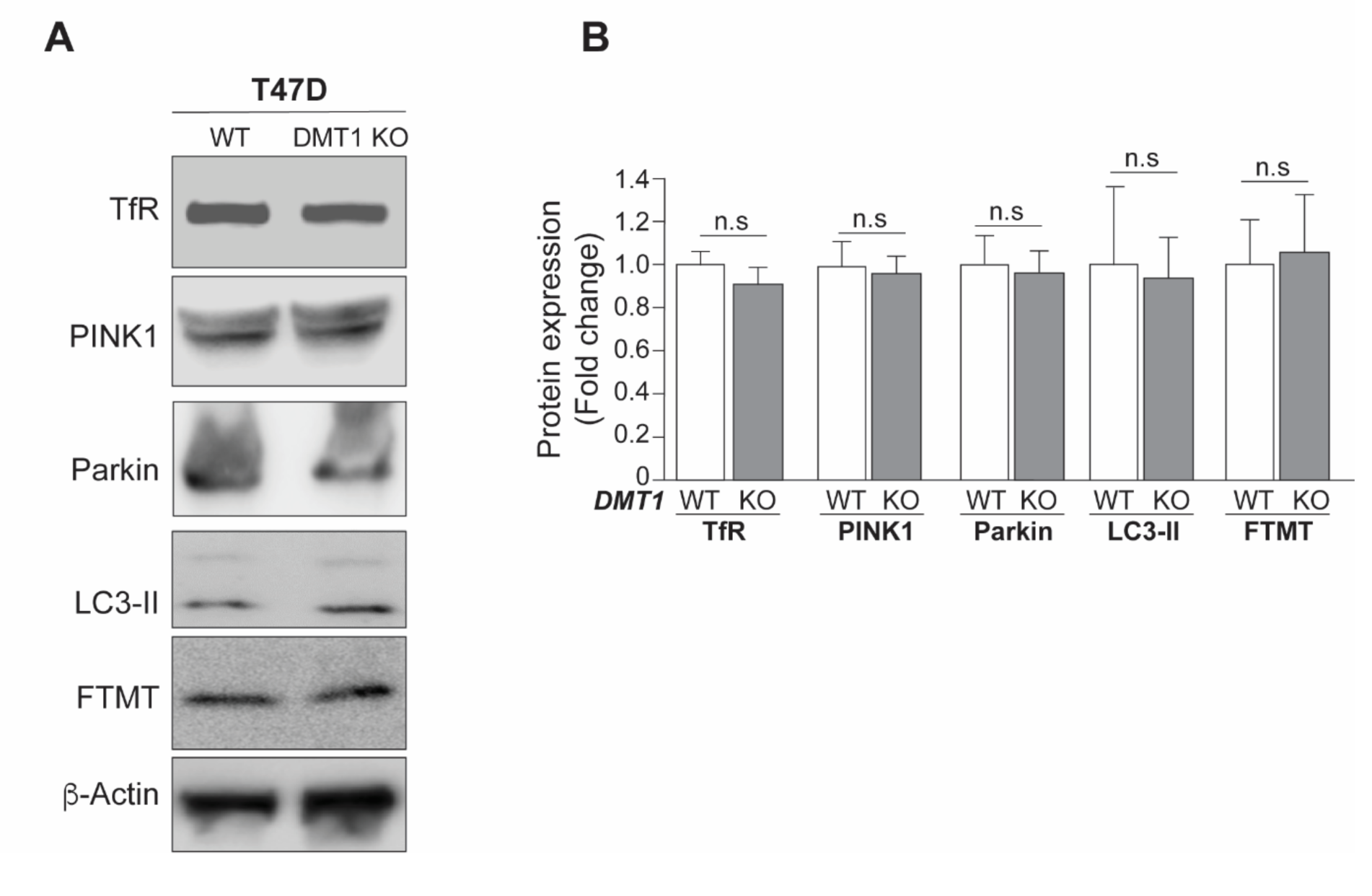
In T47D breast cancer cells DMT1 silencing did not alter the protein expression levels of TfR or mitophagy related markers. **(A)** Immunoblots and **(B)** normalized densitometry quantification of TfR, PINK1, Parkin, LC3B and mitochondrial ferritin (FTMT) in T47D WT and T47D DMT1 KO cells. ²-Actin was used as a loading control. Unpaired *t*-test; n = 3; ns = not significant.

**Figure Supplementary S2.**
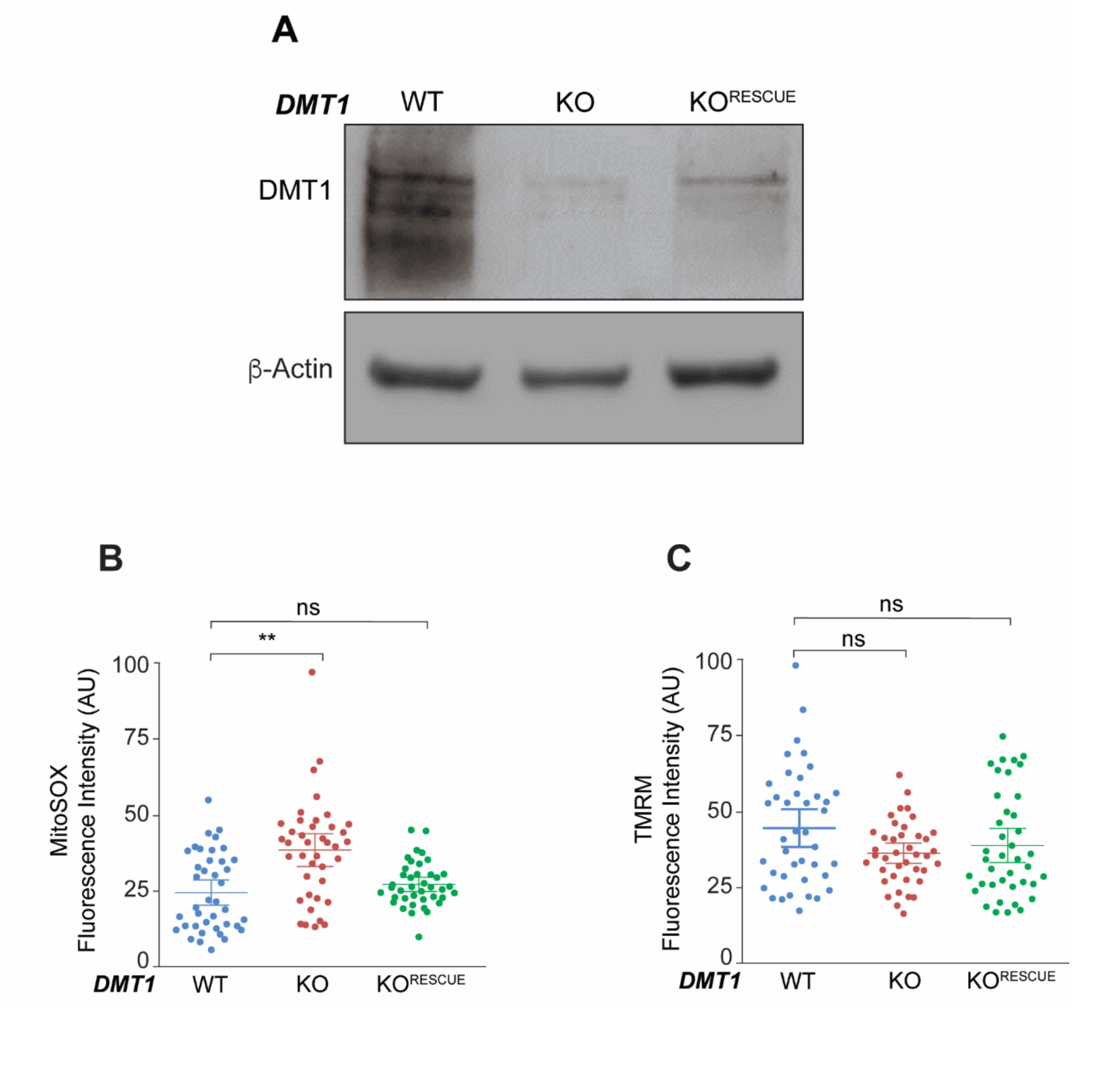
DMT1 rescue and effect of DMT1 silencing in mitochondrial superoxide production and mitochondrial membrane potential in MDA-MB-231 cells. **(A)** Representative immunoblots of DMT1 in MDA-MB-231 WT, DMT1 KO, and DMT1 KO^RESCUE^ cells. ²-Actin was used as a loading control. Normalized fluorescence quantification of MitoSOX **(B)** and TMRM **(C)** dyes in indicated cells at steady state. Graphs shows quantification of fluorescence intensity in arbitrary units (AU). Fluorescence intensity was analyzed using ImageJ software in 40 cells per condition. One-way ANOVA with Bonferroni post-hoc test. **p<0.01. ns: non-significant (p>0.05).

**Figure Supplementary S3.**
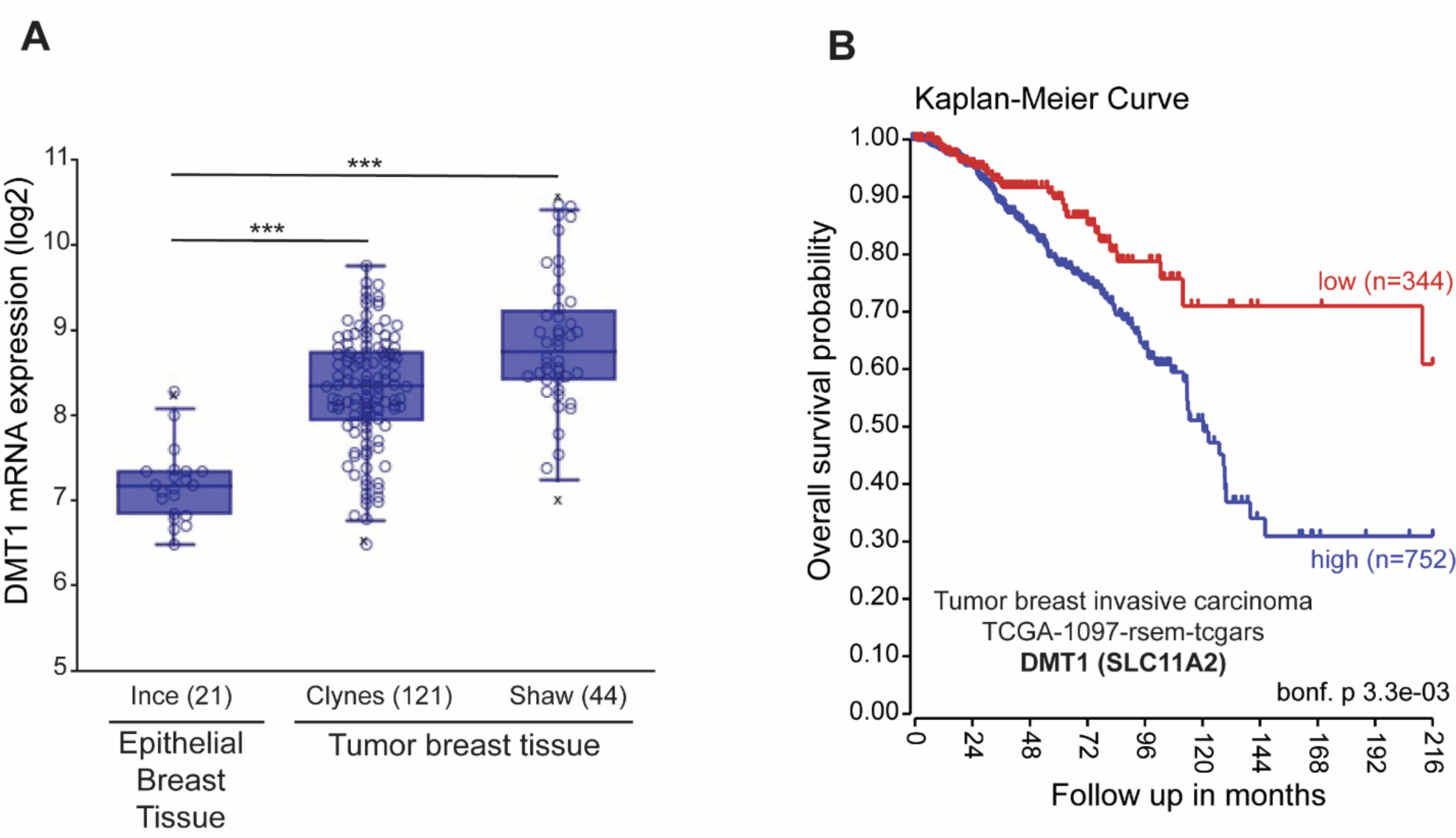
High expression of DMT1 (SLC11A2) in patient breast tumoral tissue, compared to normal breast epithelium, which is associated with lower overall survival. **(A)** 21 samples of epithelial breast tissue were compared with two breast tumor cohorts (Clynes and Shaw, 121 and 42 samples respectively) using R2 genomics analysis and visualization platform. **(B)** High DMT1 (SLC11A2) mRNA expression is associated with decreased overall survival probability in tumor breast invasive carcinoma patients. Kaplan Meier overall survival probability analysis of a cohort of tumor breast invasive carcinoma patients is shown. Long rank test result is indicated. The cut-off was calculated using the scanning method.

**Figure Supplementary S4.**
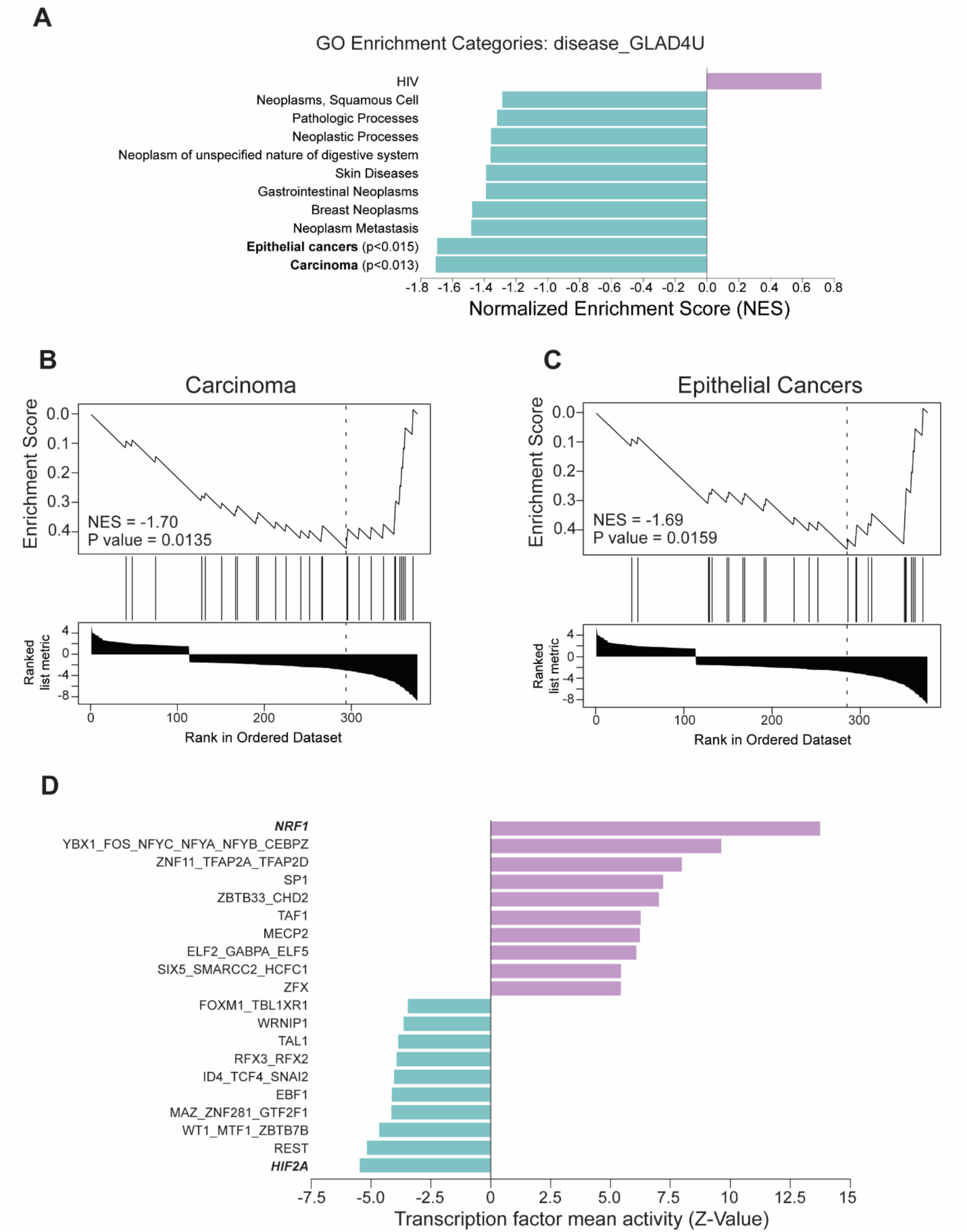
Gene Ontology (diseases) and transcription factor activity analyses from RNA seq data upon DMT1 KO in MDA-MB-231 cells. **(A)** GO analysis expressed as normalized enrichment score (NES) of diseases (GLAD4U database) using WebGestalt online platform (www.webgestalt.org). Gene Set Enrichment Analysis plots for statistically significant enrichment terms “Carcinoma” **(B)** and “Epithelial Cancers” **(C)** are depicted. **(D)** ISMARA analysis showing specific increase (purple) and decrease (green) transcription factor activity (Z-value). Nuclear factor erythroid 2-like 1 (*NRF1*) and hypoxia inducible factor 2A (*HIF2A*) are highlighted as the top increase and top decrease in transcription factor mean activity, respectively.

**Table S1.** List of primary and secondary antibodies.

| **Table S1. List of primary and secondary antibodies** |  |  |  |
| --- | --- | --- | --- |
| **Antibodies** | **Name** | **Catalog number** | **Manufacturer** |
| **Primary** | Mouse anti-SLC11A2 (DMT1) | ab55735 | Abcam |
|  | Rabbit anti-TOM20 | ab186735 | Abcam |
|  | Mouse anti-TOM20 | SC-17764 | Santa Cruz Biotechnology |
|  | Rabbit anti-PMPCB | 16064-1-AP | Proteintech |
|  | Mouse anti-Transferrin Receptor | ab1086 | Abcam |
|  | Rabbit anti-Ferroportin (SLC40A1) | NBP1-21502SS | Novusbio |
|  | Rabbit anti-PINK1 | 6946S | Cell Signaling |
|  | Mouse anti-Parkin | 4211S | Cell Signaling |
|  | Rabbit anti-mitochondrial ferritin | ab124889 | Abcam |
|  | Rabbit anti-LC3B | 3868S | Cell Signaling |
|  | Mouse anti-beta-Actin-Peroxidase | A3854 | Millipore-Sigma |
| **Secondary** | Donkey anti-mouse Alexa Fluor 488 | ab181289 | Abcam |
|  | Donkey anti-rabbit Alexa Fluor 568 | ab175694 | Abcam |
|  | Rabbit HRP-conjugated IgG | 7074S | Cell Signaling |
|  | Mouse HRP-conjugated IgG | 7076S | Cell Signaling |

**Table S2.** Chemicals/Reagents.

| **Table S2. Chemicals/Reagents** |  |  |
| --- | --- | --- |
| **Name** | **Catalog number** | **Manufacturer** |
| Protease inhibitor cocktail | 04693116001 | Roche |
| 4X Laemmli sample buffer | 1610747 | BioRad |
| 4-20% Mini-Protean TGX gels | 4561094 | BioRad |
| Bovine Serum Albumin | A9647 | Millipore-Sigma |
| 2-b-Mercaptoethanol | M3148 | Millipore-Sigma |
| Clarity Max Western ECL | 1705062 | BioRad |
| DMEM high glucose, no glutamine, no phenol red | 31053-028 | ThermoFisher |
| Fetal Bovine serum (FBS) | 30-2020 | ATCC |
| Fish gelatin solution (FGS) | G7765 | Millipore-Sigma |
| DAPI | D1306 | ThermoFisher |
| Mitotracker Red FM | M22425 | ThermoFisher |
| FerroOrange | F374 | Dojindo |
| MitoSOX Red | M36008 | ThermoFisher |
| TMRM | T668 | ThermoFisher |
| Transferrin Alexa Fluor-488 | T13342 | ThermoFisher |
| Transferrin Alexa Fluor-647 |  | ThermoFisher |
| Sodium Piruvate (100 mM) | 11360070 | ThermoFisher |
| D-(+)-Glucose solution | G8769 | Millipore-Sigma |
| Matrigel® Matrix | 356237 | BD Biosciences |
| Puromycin Dihydrochloride | A1113803 | ThermoFisher |
| Geneticin (G418 Sulfate) | 11811023 | ThermoFisher |
| TransIT®-2020 Transfection Reagent | MIR 5400 | Mirus |
| Polybrene Infection/Transfection Reagent | TR-1003-G | Millipore-Sigma |

**Table S3.**
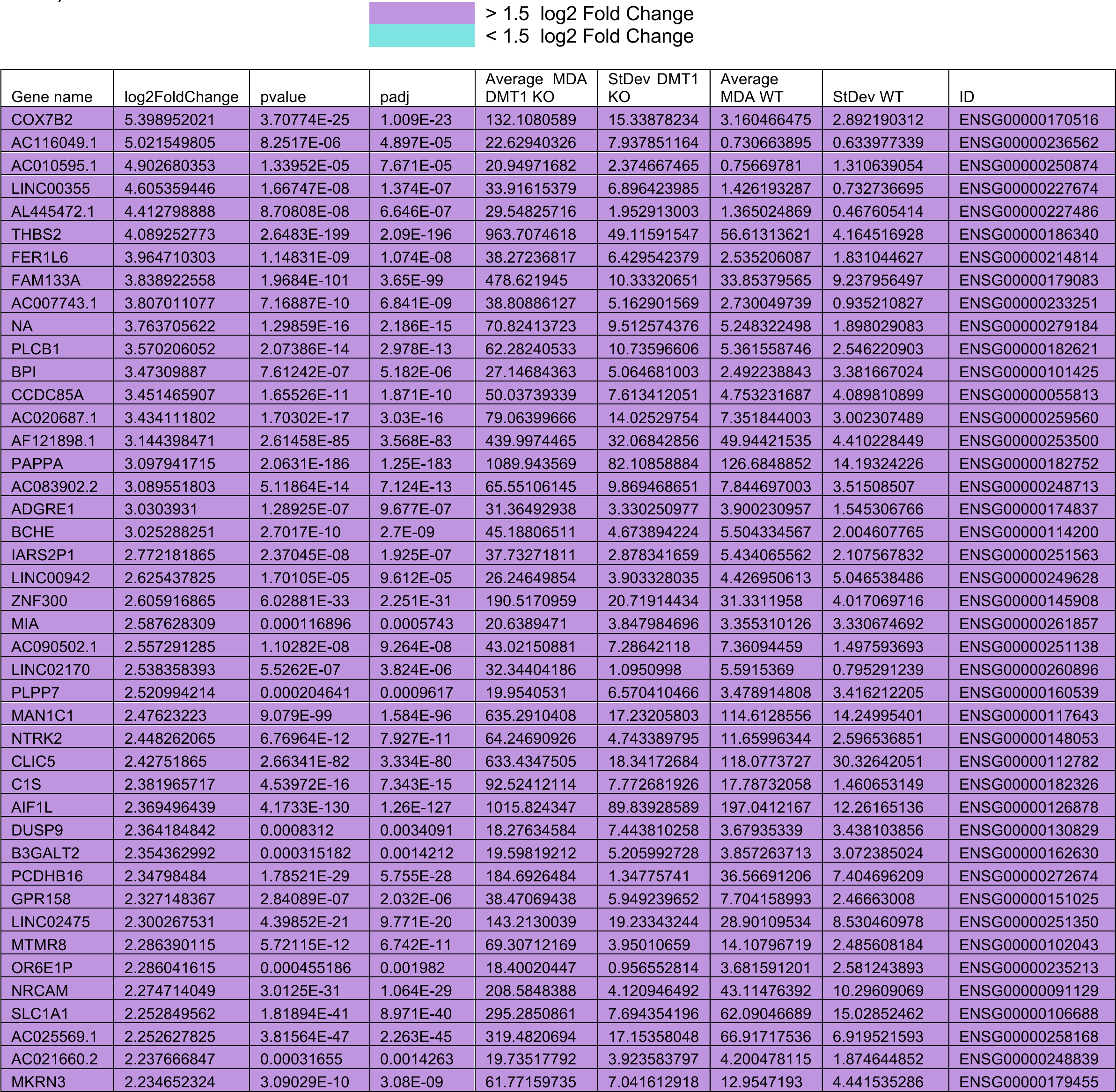
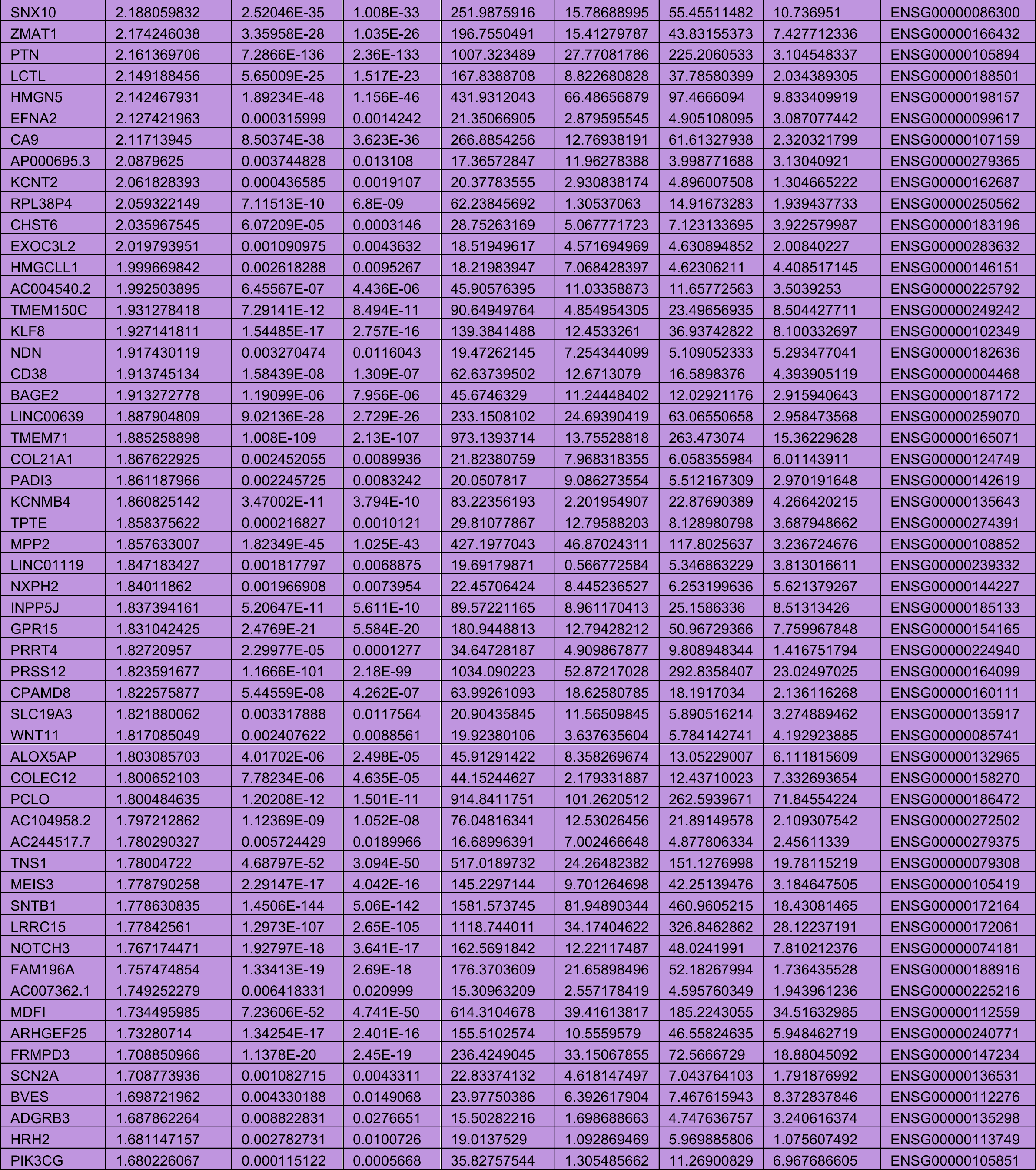
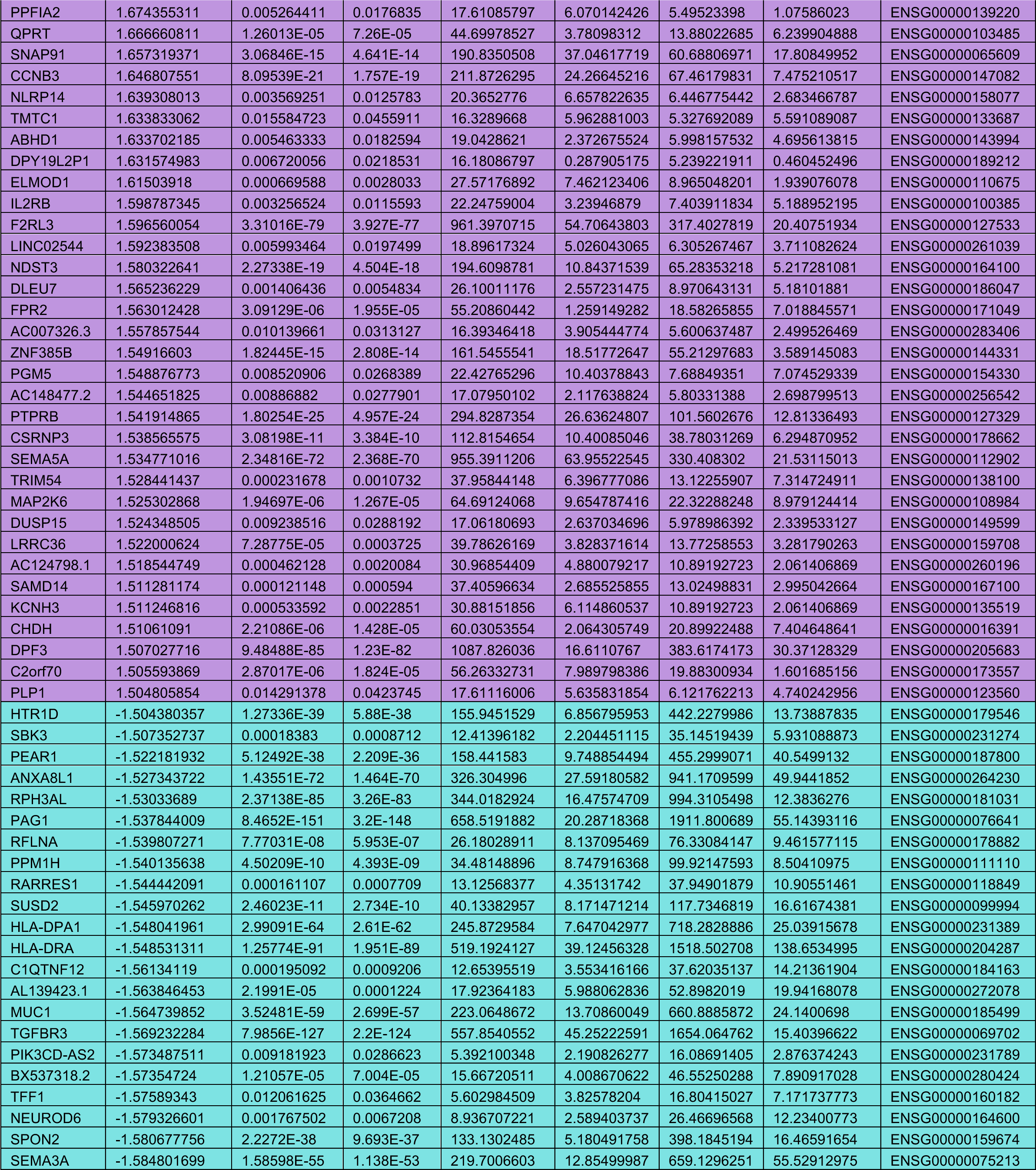
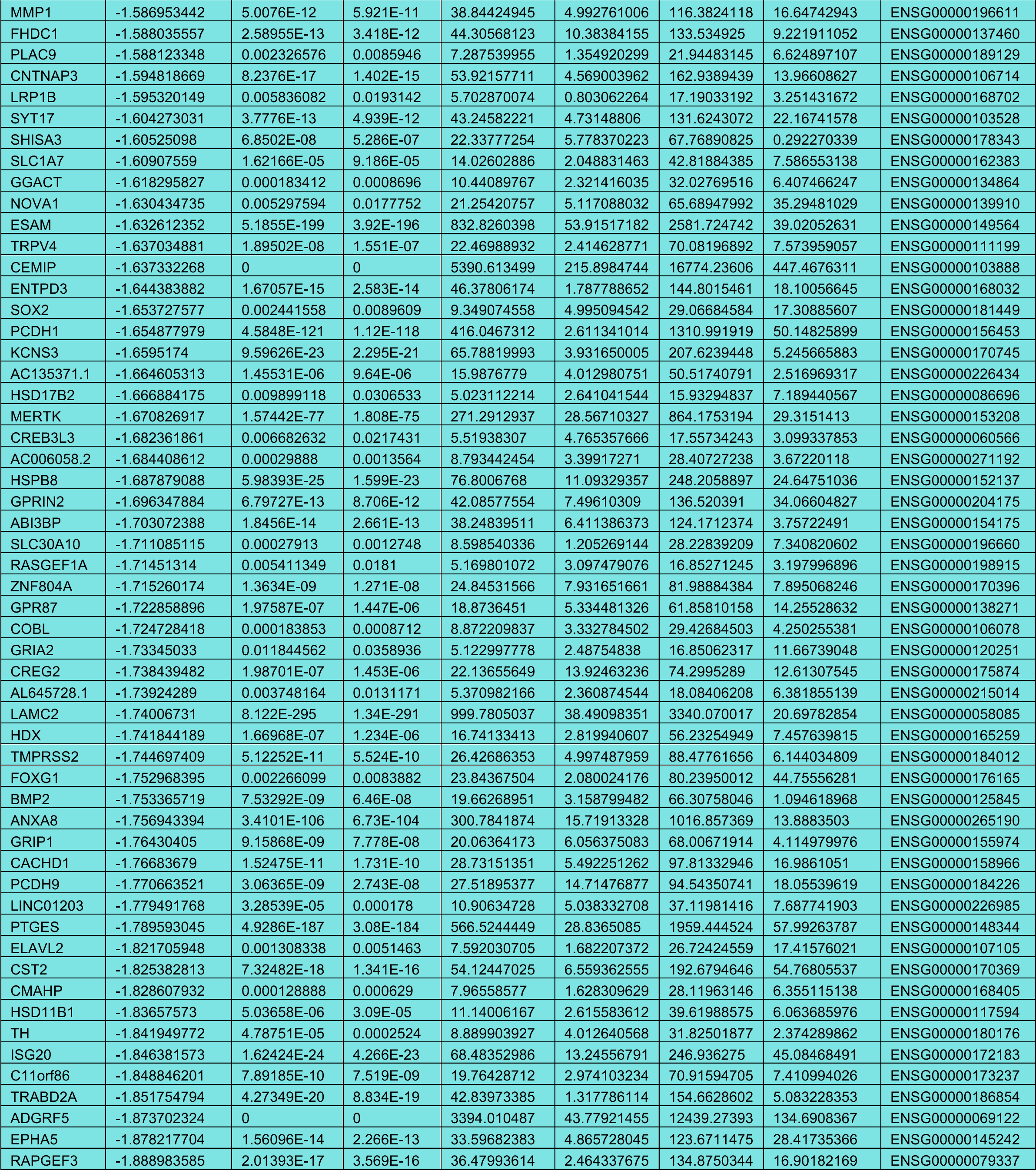
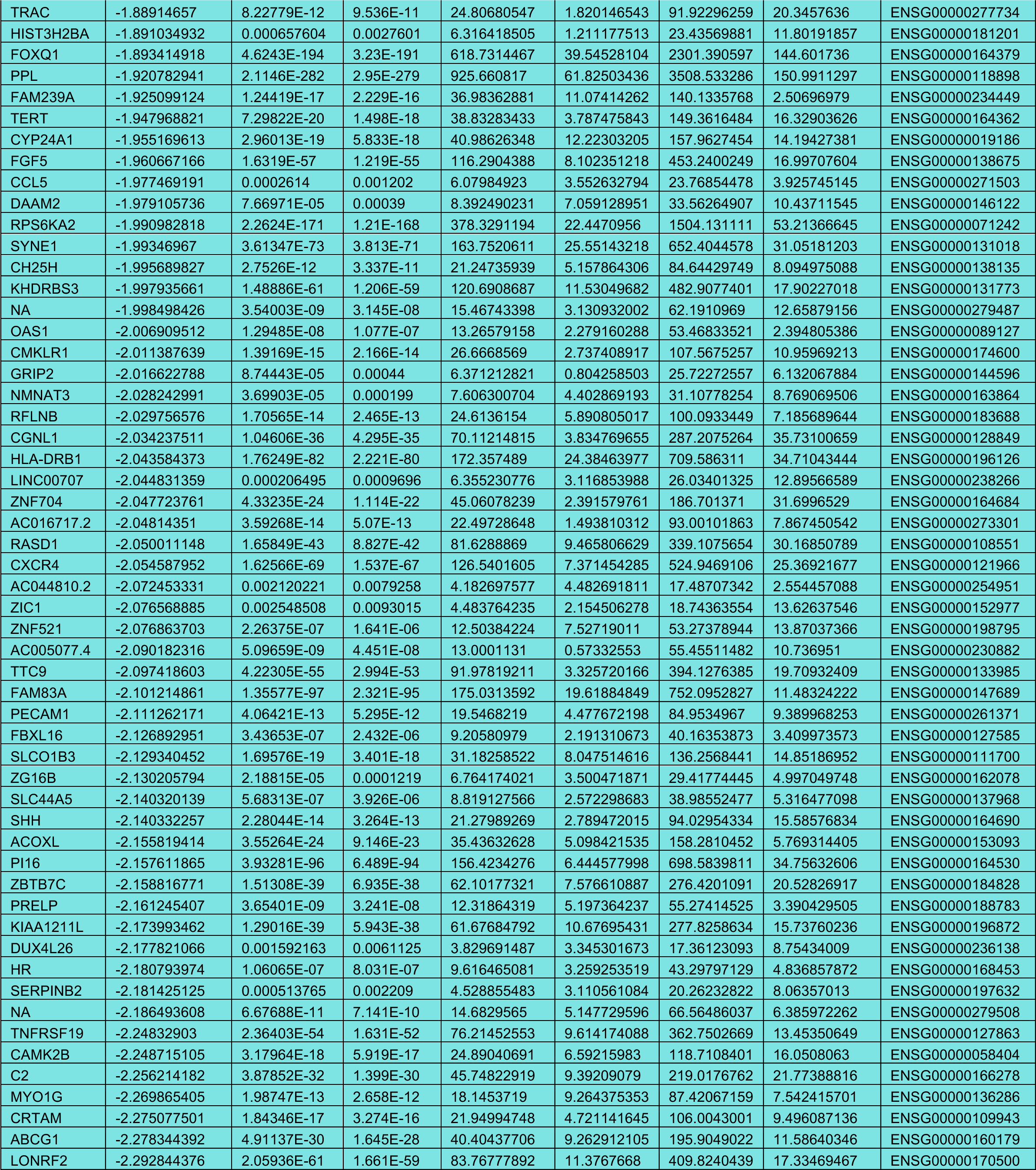
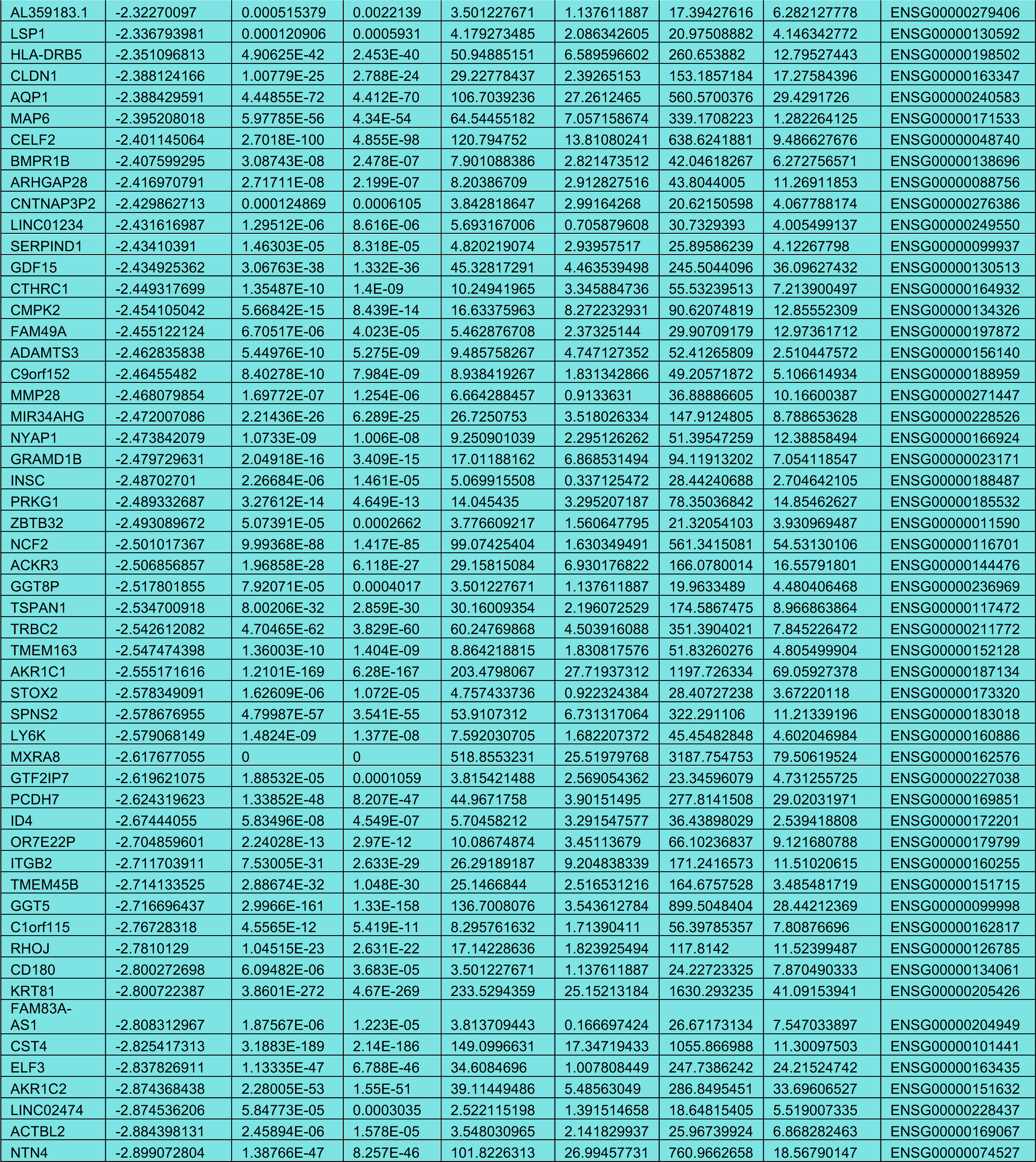
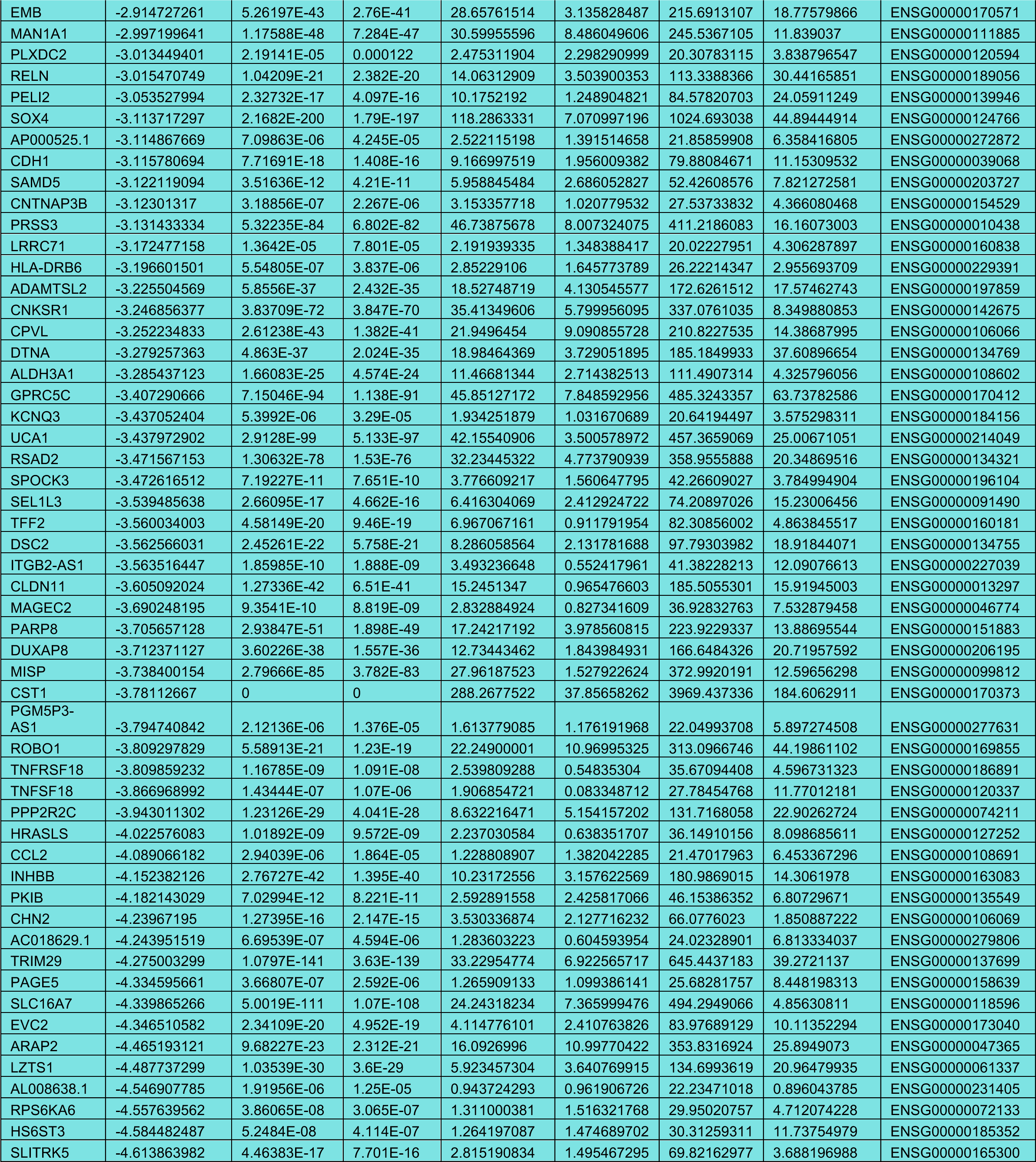
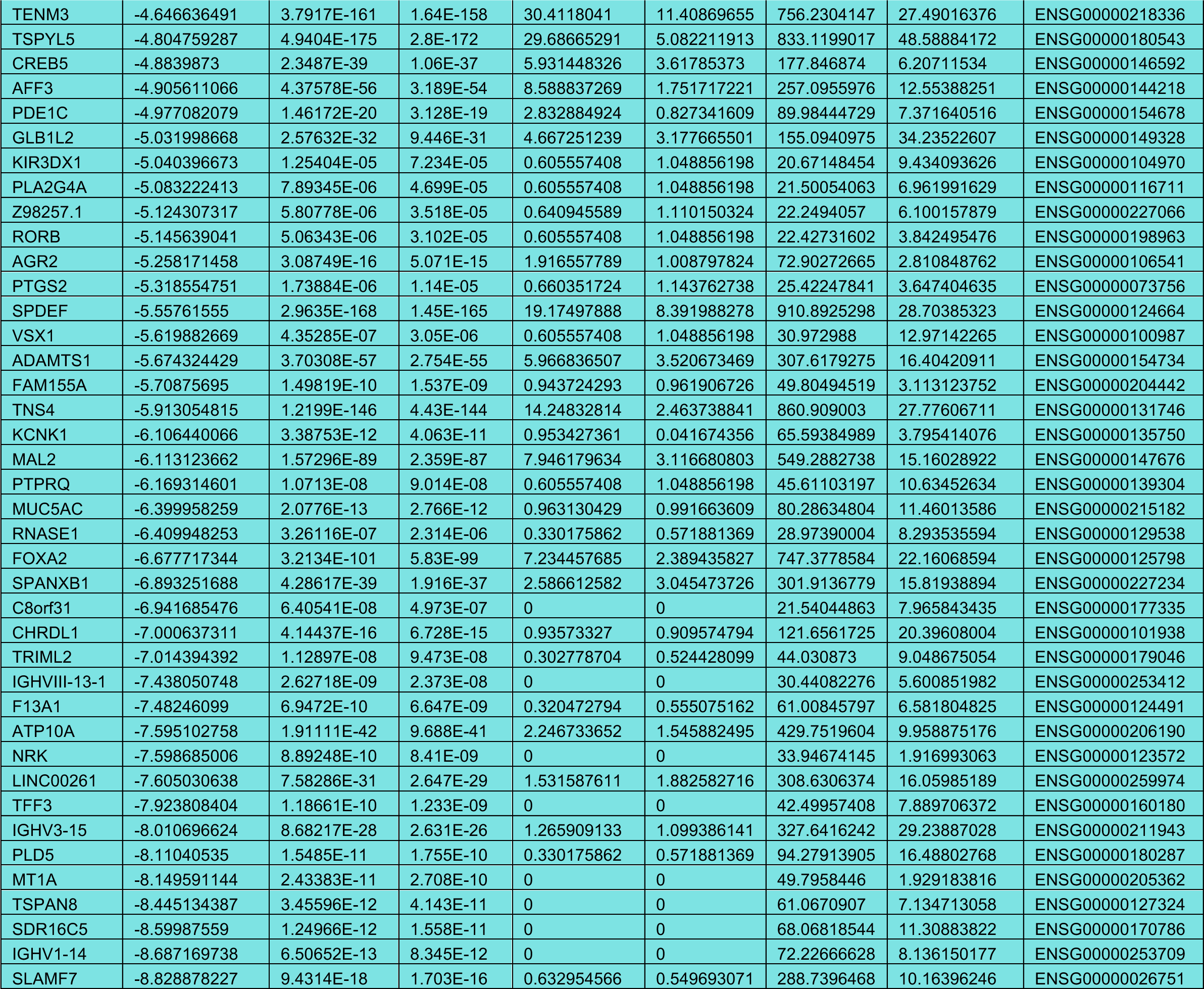
Differentially expressed genes upon DMT1 KO in MDA-MB-231 cells (RNA sequencing data).

